# Habitat and complex life cycles promote morphological diversity in salamander limb bones

**DOI:** 10.1101/2025.10.20.683565

**Authors:** Jonathan M. Huie, R. Alexander Pyron, Sandy M. Kawano

## Abstract

Salamander evolution featured multiple transitions between water and land that likely promoted distinct adaptations in limb bones for buoyancy control versus increased load-bearing capacities, respectively. Many extant species spend their entire lives either in water or on land, while, others undergo water-land transitions within their lifetime. However, exposure to both environments may impose competing demands that restrict adaptive evolution for a particular habitat. Using a 3D morphological dataset of 133 species spanning the phylogenetic and ecological breadth of salamanders, we find that the external and internal morphology of limb bones are evolutionarily decoupled, which increases the evolvability of limb bones in response to diverse mechanical demands. Terrestrial salamanders have stiffer bones with greater resistance to fracture, while aquatic species have denser bones that are hypothesized to aid in buoyancy regulation. We uncover a functional trade-off between stiffness and density that promotes stiff yet lightweight bones in terrestrial lineages. Released from terrestrial constraints, aquatic paedomorphs have disparate external morphologies, whereas terrestrial direct developers consistently share a rod-like bone shape. Aquatic and terrestrial multiphasic taxa show less morphological divergence than monophasic species living in comparable habitats but are not constrained by their complex life cycle. Multiphasic species have distinct external limb bone shapes that have evolved as fast or faster than monophasic species. Taken together, we propose that the trade-offs imposed by different habitats and complex life cycles increase limb bone diversity by promoting alternate evolutionary pathways.

## INTRODUCTION

Habitat transitions expose animals to new physical demands that often promote adaptive changes to the musculoskeletal system (Easterling et al. 2022; Koehl 2023). However, many animals have complex life cycles, which involve ecological shifts over ontogeny that expose them to conflicting selective pressures (Hanken 1992; Laudet 2011; McMenamin and Parichy 2013; Truman 2019). While metamorphosis potentially decouples selective pressures across life stages (Ebenman 1992; Hanken 1992; Moran 1994), complex life cycles may limit overall adaptive responses if the different life stages occupy habitats with dramatically different demands. For example, water-land transitions involve dramatic ecological shifts with profound consequences for the anatomy and function of limbs (Bejder and Hall 2002; Pierce et al. 2020; Dickson et al. 2021; Molnar et al. 2021a). Compared to terrestrial habitats, limbs used underwater experience lower mechanical loads due to positive buoyancy and the use of locomotor modes that minimize limb-substrate interactions, such as axial-based swimming (Azizi and Horton 2004; Young and Blob 2015; Gamel et al. 2024). In contrast, terrestrial locomotion requires the limbs to support body weight against gravity and generate larger propulsive forces than aquatic locomotion that can exceed body mass by orders of magnitude (Biewener and Taylor 1986; Biewener and Patek 2018). Consequently, the move onto land is associated with the evolution of stronger limb bones to avoid fracture (Dickson et al. 2021). However, we currently lack a clear understanding of whether complex life cycles, that expose individuals to water and land, may constrain biomechanical adaptations during evolutionary transitions.

Salamander (Amphibia: Urodela) evolution is characterized by repeated water-land and life cycle transitions, making them an advantageous system to investigate the factors that affect morphological evolution in the limbs (Baken and Adams 2019; Bonett et al. 2022). Many salamanders have a two- or three-part life cycle (multiphasic), that begins with aquatic larvae that subsequently metamorphose into adults with varying degrees of terrestriality (Bonett et al. 2022). However, even terrestrial multiphasic species return to the water for reproduction. Other lineages have monophasic life cycles and either forego metamorphosis (paedomorphism) or undergo metamorphosis in ovum (direct development), which are associated with obligate aquatic or terrestrial habits, respectively (Bonett et al. 2022). Free from contrasting environmental demands during ontogeny, monophasic salamanders may have limb bones more finely tuned to their respective environments and exhibit greater ecomorphological divergence than multiphasic species.

The external morphology of limb bones is often associated with habitat use, but the internal morphology may reflect mechanical adaptations that cannot be gleaned from external morphology alone (Kilbourne and Hutchinson 2019; Keeffe and Blackburn 2022; Rickman et al. 2023; Vera et al. 2020, 2025). Externally, long and gracile bones facilitate speed and maneuverability while short robust bones are associated with strength and increased force production (Polly and Hall 2007). Internally, compact or solid (‘dense’) limb bones with thickened cortical walls are commonly found in aquatic tetrapods; presumably for buoyancy regulation (Wall 1983; Canoville and Laurin 2010; Houssaye 2013). While denser limb bones are more fortified, they are also heavier, which can reduce mobility on land. Alternatively, terrestrial species might have evolved bones that are less dense and therefore more lightweight, but with cross-sectional morphologies that still increase stiffness (Currey and Alexander 1985). The second moment of area, a proxy for ‘stiffness’ measured from cross-sectional images, reflects a bone’s ability to resist applied forces, such as those experienced during locomotion (Lieberman et al. 2004; Huie et al. 2022). Stiffer morphologies typically entail hollower bones and wider cross-sections that distribute material further from the centroid, with the caveat that they must be thick enough to avoid buckling and fracturing. Contrasting demands between lightweight versus dense bones likely impose a trade-off between stiffness and density that is more pronounced in terrestrial lineages compared to aquatic species due to the larger effects of gravity on land. A question that then arises is how lineages respond to these opposing demands while also transitioning between conflicting life stages.

A potential strategy for mitigating the consequences of functional trade-offs is to decouple locomotor structures. For instance, weaker evolutionary integration between external and internal limb bone morphologies would allow them to evolve independently when exposed to different selective pressures (Rickman et al. 2023). Similarly, decoupling of the forelimb and hindlimb bones would enable them to take on more diverse functional roles (Gatesy and Dial 1996; Vera et al. 2025). During most limb-based locomotion in salamanders, the hindlimbs serve as the primary propulsors and the forelimbs mainly act as brakes (Kawano et al. 2015; Dickson et al. 2021; Kawano and Blob 2022). The external forces applied to the limbs of the aquatic newt *Pleurodeles waltl* during terrestrial locomotion are lower and more divergent between their forelimbs and hindlimbs compared to the terrestrial salamander *Ambystoma tigrinum* (Kawano and Blob 2022). Additionally, the demands of terrestrial locomotion are thought to constrain limb length and maintain stronger covariance between the limbs of terrestrial salamanders compared to aquatic lineages (Ledbetter and Bonett 2019). Thus, hindlimb bones may be substantially stiffer than forelimb bones in more aquatic lineages. Yet, terrestrial salamanders interact with their environment in diverse ways (i.e., climbing, digging, jumping) (Blankers et al. 2012; Baken and Adams 2019) that may promote greater functional decoupling between their limbs to accommodate different loading regimes. Variation in the developmental timing of forelimbs and hindlimbs may also influence patterns of limb integration. Many paedomorphs and multiphasic species develop their limbs disjointly, while direct developers hatch with all four limbs already developed (Bonett and Ledbetter 2022), suggesting that direct developers may exhibit stronger levels of covariance between limbs than other life cycle strategies.

We examined the effects of habitat preference, life cycle strategy, and functional trade-offs on the external and internal morphology of limb bones using micro-computed tomography (micro-CT) scans of 133 species spanning the phylogenetic and ecological breadth of salamanders. We used 3D geometric morphometrics to compare external shape, cross-sectional geometries to compare density and second moment of area (‘stiffness’), and phylogenetic comparative methods to quantify the tempo and mode of evolutionary patterns in the stylopods (i.e., humerus and femur). We hypothesized that terrestrial salamanders have stiff yet lightweight limb bones, represented by a functional trade-off (i.e., strong negative correlation) between stiffness and density. Without the constraints of terrestrial locomotion, aquatic species may exhibit greater morphological disparity and a functional trade-off that is weaker or absent. We also hypothesized that multiphasic species would exhibit less morphological divergence than their monophasic counterparts due to exposure to ontogenetic habitat transitions in the former. Finally, we hypothesized that the mechanical requirements of terrestrial locomotion promoted stronger integration between forelimbs and hindlimbs. Our work enhances our understanding of how biomechanical factors and contrasting ontogenetic demands influence the evolutionary changes of locomotor structures across water-land transitions.

## METHODS

### Morphological and taxon sampling

We examined the humeri and femora of 133 salamander species, including representatives from all 10 families and 65 of the 69 (94%) extant genera (AmphibiaWeb 2025). Each species was represented by a single specimen due to the limited availability of public CT scans. All specimens were considered adults or large juveniles based on their body size. Most scans were obtained from MorphoSource.org (Boyer et al. 2016; Blackburn et al. 2024), but 23 additional scans were generated for this study. Scanning was conducted at the Friday Harbor Labs Karel F. Liem Bio-Imaging Center, using a Bruker Skyscan 1173, or at the University of Michigan Museum of Zoology (UMMZ) Micro(µ)CT Scanning Laboratory, using a Nikon XT H 225ST. One humerus and one femur per specimen were selected for morphological analysis, except for sirens (Sirenidae) that have secondarily lost their hindlimbs. To increase data quality, all sampled limb bones were required to be at least 100 slices long. We used 3D Slicer (version 5.6.1) with the SlicerMorph toolkit (version 23d21fd) to visualize, isolate, and generate models of each limb bone (Kikinis et al. 2014; Rolfe et al. 2021). Bones selected from the left side of the body were mirrored so they appeared to originate from the right side.

### Ancestral reconstruction of habitat preference and life cycle strategy

Salamanders were classified based on their habitat preferences (i.e., aquatic, semi-aquatic, or terrestrial) and life cycle strategy (i.e., paedomorphic, multiphasic, direct developer). Most classifications were curated from previous studies or repositories (Fabre et al. 2020; Louppe et al. 2025; AmphibiaWeb 2025), and modified as needed to more accurately reflect the natural history of each taxon. We based habitat preferences on where adults spend most of their time, with the semi-aquatic group being represented by species that live at - and regularly move across - the interface of aquatic and terrestrial environments. Newts with a multi-year terrestrial ‘eft’ stage but an aquatic adult stage or long aquatic breeding season were also considered semi-aquatic. While some multiphasic species exhibit facultative paedomorphism, we primarily sampled metamorphosed adults. In these cases, habitat and life cycle classifications were coded based on the characteristics of the sampled individuals (Table S1). Viviparous salamanders were not included in this study due to small sample sizes. A caveat with our classification approach is that salamanders are unlikely to match these discrete categories perfectly, especially semi-aquatic taxa, but they allow us to test general hypotheses surrounding habitat use and life cycle strategy.

We reconstructed the joint evolutionary history of salamander habitat and life cycle strategy using the most comprehensive time-calibrated salamander phylogeny to date (Stewart and Wiens 2025). We pruned the tree to include only species in our taxonomic sampling. Species sampled for morphology but absent from the phylogeny (*Bolitoglossa diminuta, Hypselotriton wolterstorffi*, and *Plethodon grobmani*) were added by replacing the tips of taxa in the same species complex (*Bolitoglossa aureogularis, Hypselotriton yuannensis*, and *Plethodon savannah*, respectively). We used the CorHMM R package v2.8 (Beaulieu et al. 2022) to reconstruct the evolution of habitat preference and life cycle strategy with hidden Markov models (Boyko and Beaulieu 2021). We fit three transition rate models (equal rate, symmetric, and all-rates different) with and without a hidden state parameter, and all with and without dual transitions for a total of 12 models. We generated 1,000 stochastic character mappings using parameters from the best fitting model based on weighted AIC values. All analyses were performed in R version 4.0.2 (R Core Team 2023).

### External limb bone shape

We used 3D geometric morphometrics to compare the external shapes of the humeri and femora. Salamander limb bones have few identifiable landmarks; therefore, we used a pseudolandmark approach. We generated 193 pseudolandmarks on a representative humerus (*Aneides hardii*) and 190 on a representative femur (*Plethodon elongatus*) using the PseudoLMGenerator module from the SlicerMorph toolkit in 3D Slicer (Rolfe et al. 2021) (Figure S1). The pseudolandmarks were transferred from the representative templates to the remaining humeral and femoral models with the ALPACA module (Porto et al. 2021). Landmark files were imported into R, where we performed sensitivity analyses to assess whether our results were robust to the number of landmarks using the “LaSEC” (Landmark Sampling Evaluation Curve) function in the *LaMBDA* package v0.1.1 (Watanabe 2018). Then we performed generalized Procrustes Superimpositions on the humerus and femur data sets using the “gpagen” function in the *geomorph* package v4.0.8 (Baken et al. 2021). To visualize the major axes of shape variation we conducted principal component analysis (PCA) with a covariance matrix using the “gm.prcomp” function in *geomorph*.

### Internal limb bone morphology

We measured two cross-sectional traits that are proxies for bone stiffness and density. Specifically, we calculated the maximal second moment of area irrespective of any anatomical axes and bone compactness, respectively, using the *SegmentGeometry* module for 3D Slicer (Huie et al. 2022). We calculated the cross-sectional traits across the middle 10% of the limb bones’ length, where mechanical loads are predicted to be greatest (Biewener and Taylor 1986), and calculated the average values for each bone. To isolate the effects of cross-sectional shape on bone stiffness, we normalized the second moment of area of the bones with the second moment of area of solid circles with the same cross-sectional area (Huie et al. 2022). These normalized second moment of area values (referred to as “stiffness”) represent how well a bone’s shape can resist bending relative to a solid rod. Bone compactness (referred to as “density”) measures the ratio between the cross-sectional area of the cortical bone and the total cross-sectional area (area of the cortical bone and vacuities within the section). Total cross-sectional areas were calculated from solid versions of our limb models generated with the *SurfaceWrapSolidify* module in 3D Slicer (Weidert et al. 2020).

### Statistical analyses

We performed phylogenetic ANOVAs to test the effects of body size, habitat preference, life cycle strategy, and their interactions on limb bone morphology using the “procD.pgls” *geomorph* function. Due to the absence of some habitat and life cycle combinations *in situ*, we tested for an effect of ecotype (combined habitat and life cycle categories) on morphology. Separate ANOVAs were performed for each bone and trait using a similar formula: trait ∼ log(size) + ecotype + log(size) * ecotype. We used snout-vent length (SVL) as a proxy for size because locomotor forces are expected to scale with body size. For the phylogenetic ANOVAs involving the cross-sectional traits, we fit an Ornstein–Uhlenbeck (OU) error matrix because we determined that an OU model of evolution was a better fit for the data than a Brownian Motion (BM) model with the “fitContinuous” function in the *geiger* package v2.0.11 (Pennell et al. 2014). Where ecotype had a significant effect on shape (p ≤ 0.05) or there was an interaction between size and ecotype, we performed pairwise post-hoc tests to compare the mean shape differences and allometric slopes between ecotypes, respectively. Humeral analyses were performed with and without sirens but, because the results were similar (Table S3-4), we report the results that included the sirens.

We also performed phylogenetic paired t-tests to compare differences in mean stiffness and density between the humerus and femur. Separate analyses were conducted to compare the humerus and femur of each ecotype using the “phyl.pairedttest” function in the *phytools* R package v2.4 (Revell 2024).

To characterize the effect of habitat preference and life cycle on morphological diversity, we estimated the disparity of the bone shapes and cross-sectional traits with the “morphol.disparity” function in *geomorph*. We performed a separate analysis on each trait and used ecotype as the discrete variable. For humeral traits, we performed the analyses with and without the sirens.

### Correlated evolution of limb bone traits

A functional trade-off between stiffness and density would be represented by a negative correlation between the traits. To compare the evolutionary correlation between stiffness and density across the different ecotypes, we employed the *ratematrix* R package v1.2.4 (Caetano and Harmon 2017) to implement a Bayesian approach for estimating phylogenetic variance-covariance matrices (Caetano and Harmon 2019). We used the ‘*Q’* transition-rate matrix from the ancestral state reconstruction to generate a distribution of 1,000 stochastic character maps for the evolutionary history of the five ecotypes to account for uncertainty. Then we ran multiple Markov chain Monte Carlo (MCMC) in parallel for five million generations each with 25% burn-in and discarded non-convergent chains until five convergent chains remained. We used the default parameters for all other priors. We ran separate analyses for the humerus (with and without sirens) and femur data sets.

We also investigated the evolutionary integration between forelimb and hindlimb traits across ecotypes. The Bayesian *ratematrix* approach was used to compare the correlations between the humeri and femora regarding stiffness and density. To assess the evolutionary integration of the high-dimensional geometric morphometric datasets, we used the “phylo.integration” function in *geomorph* to perform a two-block partial least-squares analysis to quantify shape covariance between the external shapes of the humerus and femur. Separate analyses were run to compare the limbs within each ecotype. Sirens were omitted from all tests that quantified the integration between the forelimbs and hindlimbs.

### Evolution of limb bone shape

We tested whether habitat preference and life cycle strategy influenced the evolution of cross-sectional shapes with an evolutionary model fitting approach. We fit each trait with a set of 26 evolutionary models using the recently described *hOUwie* framework implemented in the *OUwie* R package v2.13 (Boyko and Beaulieu 2021). Parameterized model structures were used to test for both correlated (CD; character dependent) and uncorrelated (CID; character independent) evolution between the cross-sectional traits, habitat preference, and life cycle strategy. We first fit a BM model (BM1) with a single evolutionary rate parameter (σ^2^), and a single-rate and single adaptive optimum (θ) OU model (OU1). We also fit more complex models with varied rate parameters (BMV, OUV), trait optima (OUM, OUMV), or rates and trait optima (OUMV) per selective regime. We fit three versions of the BMV, OUV, OUM, and OUMV models with modified model structures to reflect three regime classifications based on habitat preference, life cycle strategy, and the combined ecotypes. Because the evolution of cross-sectional shapes may be affected by factors unaccounted for in our analyses, we also fit CID versions of the BMV, OUV, OUM, and OUMV models each with an additional rate category to account for hidden states. Each model was fitted with 100 stochastic maps per iteration, and with adaptive sampling enabled for the CID models. Model-averaged evolutionary rate and trait optima were estimated for each ecotype using AICc scores to weight the models.

There are currently no reliable methods to model the effects of a discrete character, much less the joint effect of two characters, on high-dimensional data in an OU framework (Adams and Collyer 2019). Therefore, we used a multivariate Brownian Motion framework implemented in *geomorph* to estimate differences in evolutionary rates for external limb bone shape. We used the five ecotype categories as the discrete variable and estimated humeral rates with and without sirens.

## RESULTS

### Evolutionary history of habitat and life cycle

Evolutionary changes in habitat and life cycle were best described by symmetric transition rates and simultaneous changes (Figure 1). The ancestor of modern salamanders had a high probability of being either an aquatic paedomorph (47%) or terrestrial multiphasic species (48%), and a small chance of being semi-aquatic and multiphasic (5%). We recovered multiple transitions and reversals between aquatic, semi-aquatic, and terrestrial habitats for multiphasic species. We also estimated up to ten independent transitions to aquatic paedomorphism from a multiphasic ancestor and nine reversals. Consistent with previous studies, we found a single transition to direct development and two or three independent reacquisitions of an aquatic larval stage (Chippindale et al. 2004; Bonett et al. 2014).

**Figure 1.**
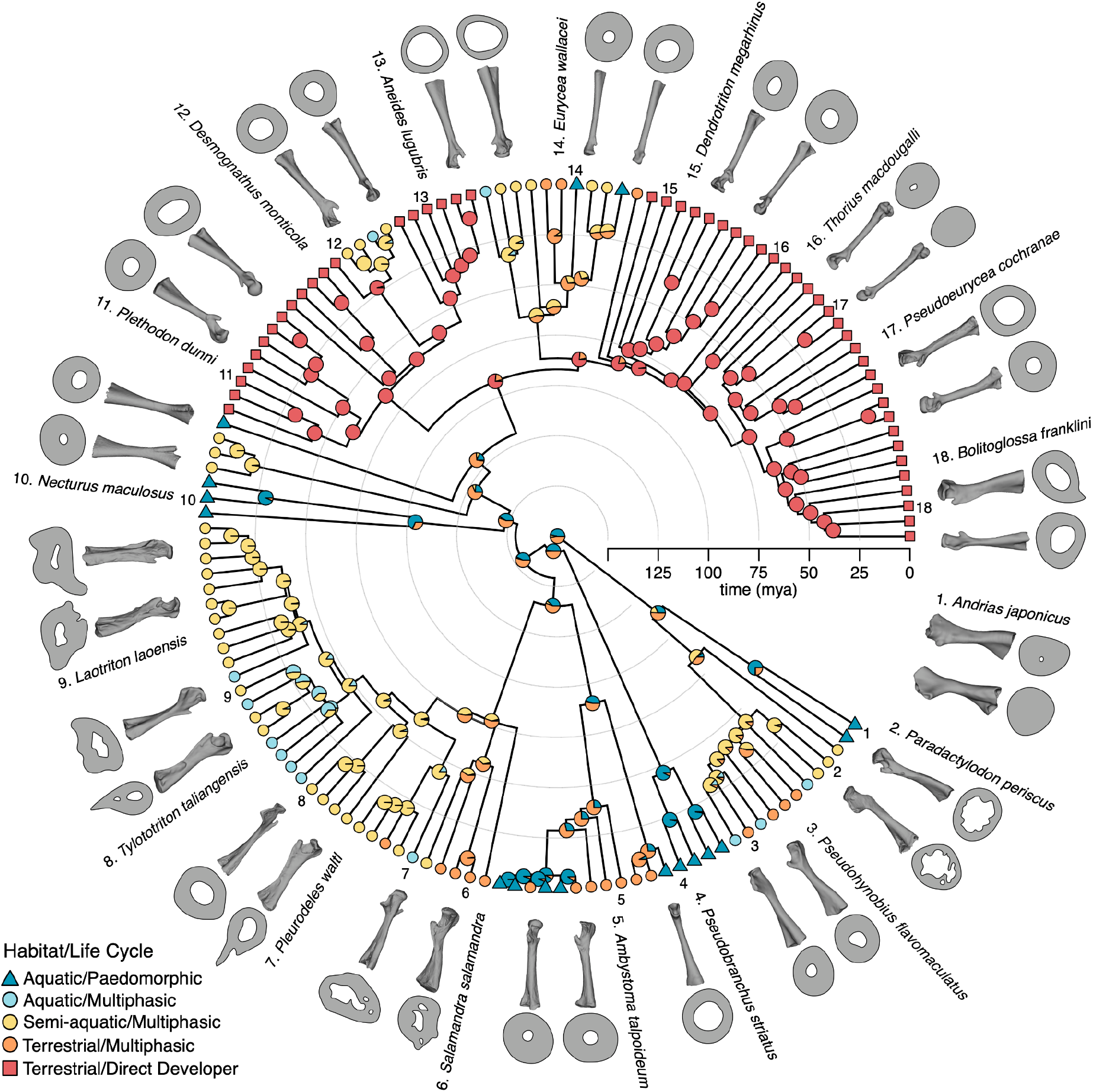
Phylogeny of 133 salamander species sampled for morphological analysis, showing the joint ancestral state reconstruction of habitat and life cycle. Insets depict representative humeri and femora and their cross-sectional shapes at the midshaft (not to scale).

### Disparity in external morphologies

Salamanders displayed a broad range of humeral and femoral shapes with similar axes of diversification (Figure 2). The primary axes of separate morphospaces for humeri and femora (principal component 1, PC 1) both depicted the contrast between gracile and robust bones, consistent with modifications for mobility versus strength, respectively (Polly and Hall 2007; Young et al. 2014). PC 2 of the humeral and femoral morphospaces captured variation in the size and presence of the ossified humeral and femoral heads. PC 3 reflected variation in the curvature of the bone shaft (diaphysis), possibly reflecting differences in the predictability of loading patterns (Bertram and Biewener 1988).

**Figure 2.**
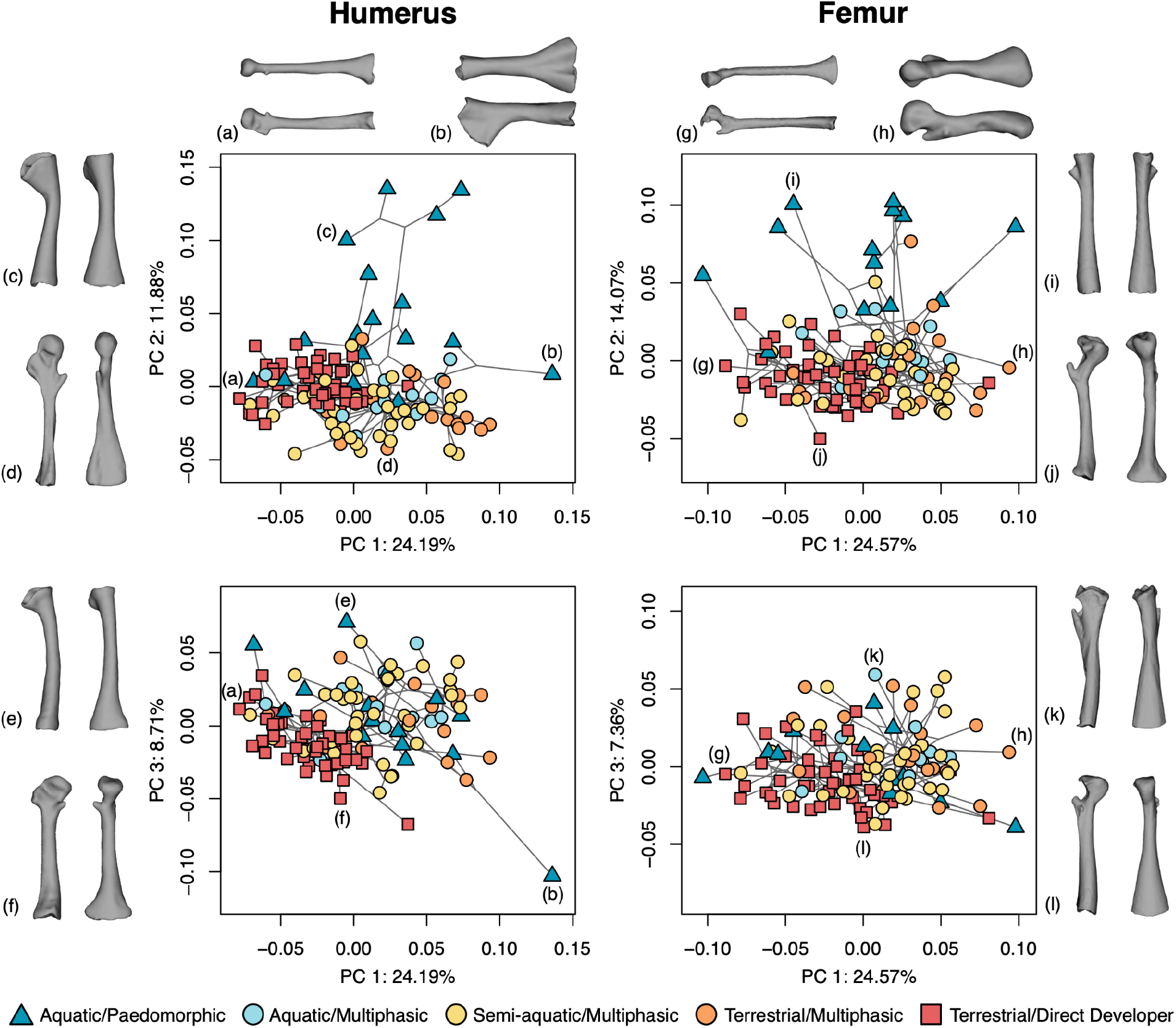
Phylomorphospaces illustrating the shape diversity of salamander humeri (left) and femora (right). Separate principal component analyses were used to identify the primary axes of variation for each limb bone. Lateral and dorsal perspectives of bones from representative taxa depict the extremes of each phylomorphospace: a) *Nyctanolis pernix*, b) *Andrias japonicus*, c) *Siren intermedia*, d) *Salamandrina terdigitata*, e) *Pseudobranchus striatus*, f) *Bolitoglossa porrasorum*, g) *Chiropterotriton magnipes*, h) *Ambystoma maculatum*, i) *Amphiuma tridactylum*, j) *Bradytriton silus*, k) *Hypselotriton wolterstorffi*, l) *Ensatina eschscholtzii*.

Terrestrial salamanders displayed lower morphological disparities in humerus and femur shapes compared to aquatic lineages (Table S2). Aquatic paedomorphs had the most diverse humeri and femora that were 2.8x (2.3x without sirens) and 1.9x more diverse than those of the terrestrial direct developers, respectively (Table S2). Multiphasic species exhibited intermediate disparities that were 0.96x-1.8x more diverse than direct developers. In both morphospaces, paedomorphs spanned the range of values along PC 1 and PC 3 and uniquely occupied a region of PC 2 associated with absent or highly reduced ossified humeral heads. Terrestrial direct developers were generally restricted to regions of the morphospaces associated with more rod-like and gracile bones. Multiphasic plethodontids also had more rod-like bones, but most multiphasic species from other families had more robust bones. All multiphasic species had larger epiphyses than aquatic paedomorphs.

Phylogenetic ANOVAs indicated that body size, ecotype, and their interactions explained significant variation in humerus and femur shape (p ≤ 0.002) (Table S3). Pairwise tests supported findings that paedomorphs had humeri and femora morphologies that were distinct from multiphasic species (p ≤ 0.015); no other comparisons were statistically significant (Table S4). Paedomorphs also exhibited distinct relationships between bone shape and body size compared to all ecotypes except for aquatic biphasic species (p ≤ 0.017) (Table S4). Larger aquatic paedomorphs had more robust bones than smaller species, whereas other ecotypes maintained relatively similar shapes at all body sizes (Figure S2).

### Disparity in internal morphologies

Terrestrial salamanders had stiffer humeral and femoral cross-sectional shapes, while aquatic salamanders had denser and more solid bones than terrestrial species (Figure 3). In contrast to patterns of external shape disparity, terrestrial salamanders exhibited more variation in their cross-sectional shapes than aquatic species (Table S2). Direct developers had cross-sectional shapes that were 2.9-8.4x (3.2-12.9x without sirens) more diverse than those of the paedomorphs (Table S2). Again, the multiphasic species were generally intermediate disparities that were 0.7-6.0x that of paedomorphs (1.2-9.2x without sirens). Within the direct developers, temperate genera (e.g., *Aneides, Hydromantes*, and *Plethodon*) generally had stiffer bones than Neotropical lineages (e.g., *Bolitoglossa, Chiropterotriton, Pseudoeurycea*). Among the semi-aquatic species, some newts (Salamandridae) and Asiatic salamanders (Hynobiidae) had the stiffest and least dense cross-sectional shapes of all species examined.

**Figure 3.**
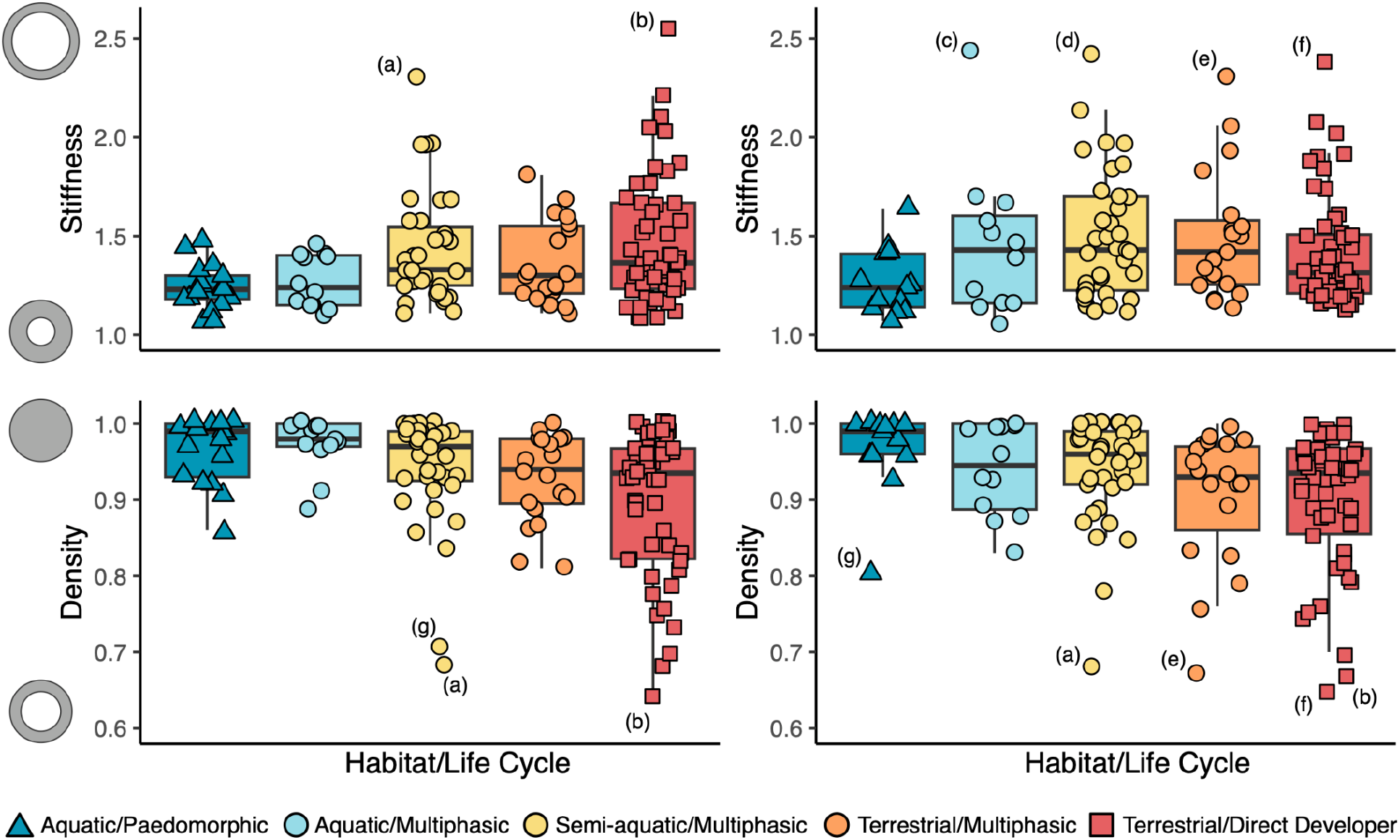
Variation in the stiffness and density of salamander humeri (left) and femora (right) across habitats and life cycle strategies. Boxplots show the median, upper and lower quartiles, interquartile range and outliers as determined by the 1.5 interquartile rule. Illustrations along the y-axes depict graphical representations of the changes in cross-sectional morphology. Some outliers are labeled: a) *Paradactylodon persicus*, b) *Aneides lugubris*, c) *Pleurodeles waltl*, d) *Tylototriton tagaliensis*, e) *Ambystoma mabeei*, f) *Hydromantes platycephalus*, g) *Onychodactylus japonicus*, and g) *Amphiuma tridactylum*.

Phylogenetic ANOVAs indicated that variation in stiffness and density is associated with differences between ecotypes (Table S3). Ecotype had a strong effect on most traits (p ≤ 0.003), except for femoral stiffness (p = 0.053). Pairwise comparisons revealed that terrestrial direct developers had significantly stiffer humeri than other ecotypes (p ≤ 0.021), except for semi-aquatic species, which also had stiffer humeri than the aquatic ecotypes (p ≤ 0.035) (Table S4). Direct developers and semi-aquatic species also had significantly stiffer femora than aquatic paedomorphs (p ≤ 0.048) (Table S4). Meanwhile, all ecotypes had significantly denser humeri than direct developers (p ≤ 0.011), but only the aquatic paedomorphs and semi-aquatic species had significantly denser femora than direct developers (p ≤ 0.010) (Table S4).

Humeral stiffness and density were also influenced by the interaction between ecotype and body size (p ≤ 0.040) (Table S3). Direct developers and semi-aquatic species exhibited steeper positive relationships between humeral stiffness and body size than aquatic paedomorphs (p ≤ 0.049), indicating that the former ecotypes have proportionally stiffer bones at larger body sizes (Figure S2). Direct developers also exhibited a steeper negative relationship between humeral density and body size than paedomorphs, terrestrial biphasic species, and semi-aquatic species (p ≤ 0.045) (Figure S2).

Despite taking on a larger propulsatory role during limb-based locomotion, the femora were not significantly stiffer than the humeri for any ecotype (p ≥ 0.230) (Table 5). Nevertheless, a few species did exhibit considerably stiffer femora than humeri, such as *Pleurodeles waltl*. There were also no differences in density (p ≥ 0.251), except for aquatic multiphasic species that had denser femora than humeri (t = 3.287, p = 0.009) (Table 5).

### Functional trade-off between stiffness and density

Stiffness and density exhibited negative evolutionary correlations, providing evidence for a functional trade-off (Figure 4A, B). Terrestrial direct developers displayed stronger median correlation values and therefore stronger trade-off between stiffness and density for both limbs (humerus/femur r = -0.880/-0.929) (Figure 4A, B). Meanwhile, aquatic paedomorphs had comparatively weaker and/or positive correlations (humerus/femur r = -0.506/0.041; when excluding siren humeri r = -0.340). However, for the humerus, aquatic (r = -0.233) and terrestrial (r = - 0.341) multiphasic species had relatively weak correlations but semi-aquatic species displayed a strong correlation (r = -0.830). For the femur, multiphasic species exhibited intermediate correlations compared to monophasic species; terrestrial (r = -0.723) and semi-aquatic (r = -0.629) multiphasic species had stronger correlations than aquatic multiphasic lineages (r = -0.192).

**Figure 4.**
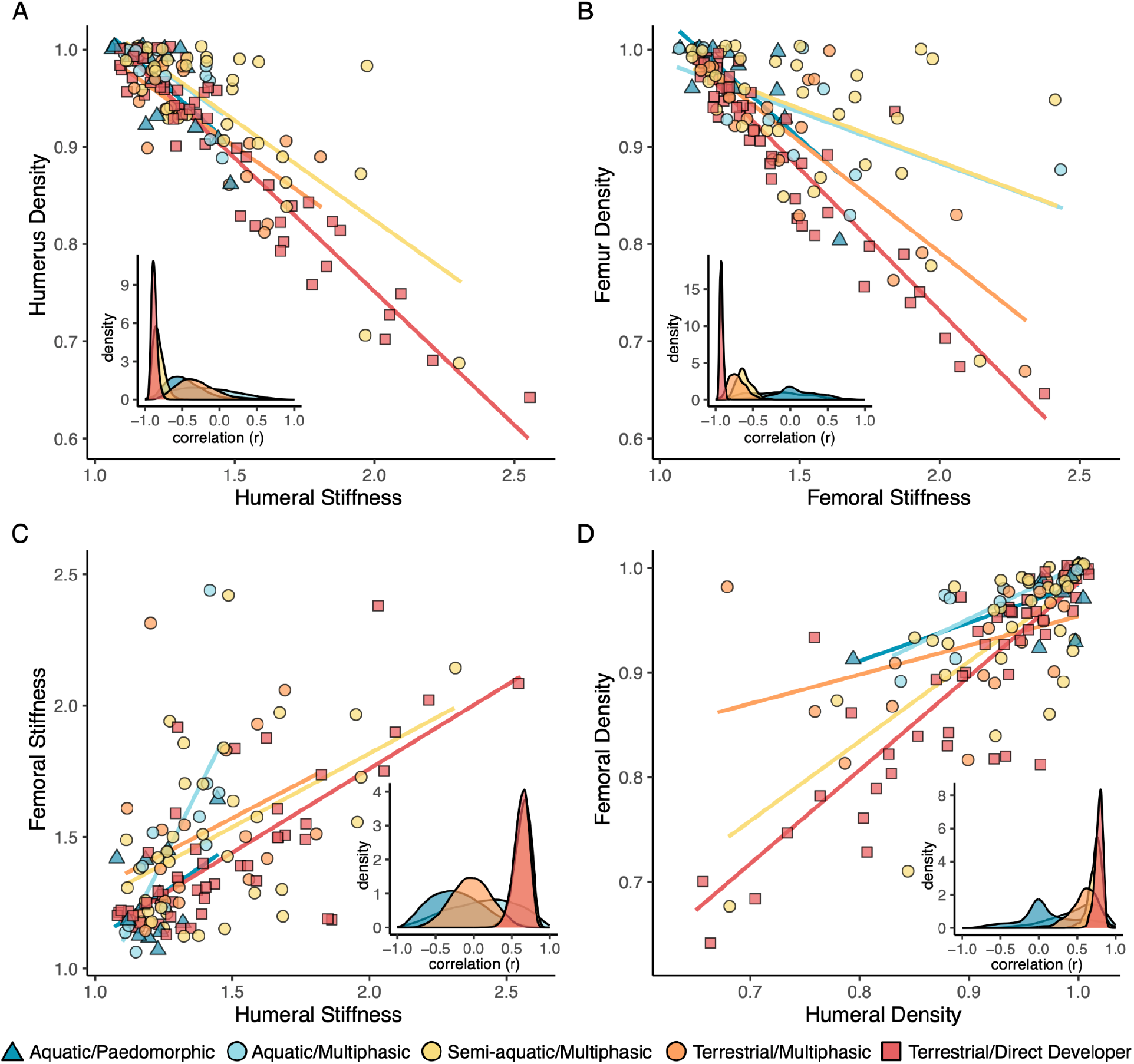
Evolutionary relationships between cross-sectional traits within and across limbs. Scatter plot points were slightly jittered to enhance readability. Insets depict density plots of the correlation (r) values estimated by Bayesian *ratematrix* analyses. Humerus plot (A) depicts the results from the analysis that included sirens. The strong negative correlations between stiffness and density within bones indicate a functional trade-off between traits (A, B). Meanwhile, strong positive correlations between the humeral and femoral traits indicate integration between the limbs (C, D).

### Evolutionary integration between limbs

Habitat preference and life cycle strategy had different effects on the evolutionary integration of external and internal morphologies between limbs. All ecotypes exhibited relatively strong signals of integration between the external shapes of their humeri and femora, based on phylogenetic two-block partial least-squares tests. Paedomorphs had the strongest integration (r-pls = 0.973, p = 0.006), followed by direct developers (r-pls = 0.929, p = 0.001), terrestrial (r-pls = 0.945, p = 0.001) and aquatic (r-pls = 0.927, p = 0.021) multiphasic species and then semi-aquatic species (r-pls = 0.882, p = 0.001). In contrast to external shape, levels of integration between the cross-sectional traits of the humeri and femora were strongest in direct-developers (stiffness/density r = 0.635/0.790) and semi-aquatic species (stiffness/density r = 0.654/0.750) (Figure 4C, D). Aquatic multiphasic species (r = 0.155) had stronger integration between humeral and femoral stiffness than terrestrial multiphasic species (r = -0.029), but the inverse was true for density (r = 0.608 and 0.563, respectively). Meanwhile, paedomorphs displayed the weakest correlations (stiffness/density r = -0.253/0.035).

### Evolutionary rates and models

Estimates of net evolutionary rates showed that the external shapes of the paedomorphic humeri and femora evolved 5.6x (6.3x without sirens) and 5.3x faster than those of terrestrial direct developers, respectively (Table S6). The limb bones of all multiphasic salamanders evolved at intermediate rates, 1.6-3.4x faster than those of the direct developers. We tested 26 models of evolution for all cross-sectional traits, and the best-fitting were generally character dependent Ornstein–Uhlenbeck models with different adaptive optima for each habitat type (OUM) or different optima and evolutionary rates (OUMV) (Tables S7–12). Model-averaged estimates of evolutionary rates indicated that the traits of the direct developers evolved 1.2–8.1x (1.2–7.9x without sirens) faster than the paedomorphs. Multiphasic species evolved at comparable rates, 0.6-9.8x faster than paedomorphs (0.8–8.3x without sirens) (Table S13). Model-averaged trait optima generally resembled the empirically measured values, with more terrestrial species associated with stiffer bones and aquatic species with denser bones (Table S13).

## DISCUSSION

Evolutionary transitions to new environments often promote functional changes to locomotor structures, but adaptations may be mediated by variation in life cycle strategies. We leveraged the rich ecomorphological diversity in salamanders to investigate the role of complex life cycles in shaping the morphological evolution of limb bones across water-land transitions. We found that aquatic and terrestrial salamanders have evolved contrasting limb bone morphologies and that multiphasic species display distinct evolutionary patterns and morphologies compared to monophasic species. Thus, we propose that complex life cycles increase, rather than constrain, diversity by promoting alternate morphological strategies capable of addressing the different demands of aquatic and terrestrial habitat use. Furthermore, external and internal regions of the limb bones have responded differently to habitat and life cycle transitions, suggesting that evolutionary decoupling provides more viable pathways for addressing functional and life history trade-offs.

### Morphological differences between ecotypes

We found evidence that terrestriality promotes stiff yet lightweight bones. Many terrestrial salamanders have limb bones with ossified epiphyses and stiffer cross-sectional morphologies that indicate they can resist greater gravitational and locomotor loads compared to aquatic lineages (Carter et al. 1998; Lieberman et al. 2004; Huie et al. 2022). Conversely, aquatic salamanders have repeatedly evolved denser limb bones, suggesting that aquatic taxa are exposed to comparable selective pressures, perhaps functioning as ballasts and increasing substrate contact during station holding or underwater walking (Canoville and Laurin 2010; Sanchez et al. 2010; Houssaye 2013; Fabbri et al. 2022). We also uncovered a functional trade-off between density and stiffness, which is more pronounced among terrestrial species. When evolving stiffer cross-sectional shapes (e.g. cross-sections with thinner walls or wider diameters), less dense limb bones also evolve in terrestrial salamanders (Figure 4A, B), presumably as a means for weight-reduction (Toyama et al. 2023). Furthermore, terrestrial direct developers have limb bones that are slender and rod-like. While large robust bones are stronger and intuitively better for dealing with the demands of terrestriality, they are also heavier and therefore more difficult and energetically expensive to move on land. These findings support that terrestriality constrains the axes of skeletal diversification in terrestrial tetrapods (Ledbetter and Bonett 2019).

Despite exhibiting narrower axes of variation, terrestrial salamanders exhibit considerable variation in cross-sectional morphology. Variation in locomotor gaits and behavior result in different magnitudes and orientations of mechanical loads being applied to the limbs that likely affect internal morphology. For instance, some scansorial salamanders (*Aneides* and *Hydromantes*) had some of the stiffest humeri and femora in our dataset, which could be adaptations for counteracting higher bone stresses and supporting a larger propulsatory role of the forelimbs during climbing (Wang et al. 2015; Munteanu et al. 2023). Moreover, salamander femora are not substantially stiffer than the humeri despite the former playing a larger role in propulsion on land, consistent with findings for the terrestrial *Ambystoma tigrinum* (Kawano et al. 2015). It is likely that non-locomotor behaviors also expose the humeri to unique mechanical demands that promote stiffer bones, such as sexually dimorphic behaviors. Males of salamandrids and hynobiids have thicker forelimbs for grabbing females during courtship (amplexus) or fighting other males (Park et al. 1996; Reinhard et al. 2015). Semi-fossorial species (*Ambystoma* spp.) also use their forelimbs for non-locomotor behaviors but did not have exceptionally stiff cross-sectional shapes expected of forelimb burrowers (Semlitsch 1983). However, other factors influence bone stiffness besides morphology, including material properties (Erickson et al. 2002; Lieberman et al. 2004). As a result, material properties may complement or mediate structural changes in response to different environmental demands.

Additionally, differences in body size also may help explain the variation in cross-sectional morphology. Because heavier animals experience disproportionately larger forces on their bones than lighter animals, the selective advantages of stiff yet lightweight bones would be greater in larger species (Currey and Alexander 1985). We found some evidence that larger animals have proportionally stiffer bones, indicated by the steeper slope between stiffness and body size among terrestrial salamanders (Figure S2). Conversely, most salamanders are relatively small animals that produce correspondingly small locomotor forces (Kawano et al. 2015; Kawano and Blob 2022). Thus, fortifying limb bones with cortical bone and increasing density may have minimal effects on mobility, especially direct developers that already have fairly gracile limb bones. That might explain why many miniature species examined herein (e.g., *Thorius macdougalli, Desmognathus wrighti, Nototriton* spp., *Parvimolge townsendi*) have relatively solid cross-sections (density values > 0.95) (Uzzell 1961; Hanken 1982). Although, *Thorius* exhibits hyper-ossification of their limb bones that coincides with sexual maturation rather than a particular body size (Hanken 1982), suggesting that not all changes in bone shape reflect mechanical adaptations for terrestrial locomotion.

### Complex life cycles promote diversity

Life cycle complexities do not influence all aspects of the salamander skeleton equally. Our results on the external shape of limb bones are consistent with previous findings that the crania and feeding apparatus of multiphasic salamanders are less morphologically diverse than those of paedomorphs, but more diverse than those of direct developers (Fabre et al. 2020; Louppe et al. 2025). Those authors proposed that complete metamorphosis, and not just the presence of a complex life cycle, constrains cranial diversity in multiphasic and direct developing species. This appears to be plausible for external limb bone shape but not for cross-sectional shape, where direct developers and multiphasic exhibit greater disparity and higher rates of morphological evolution than paedomorphs. Different from skeletal evolution, multiphasic salamanders have more constrained body shapes than monophasic species attributed to the constraints associated with conflicting environmental demands throughout ontogeny (Bonett and Blair 2017). Our findings provide additional support that complex life cycles may limit morphological diversification in multiphasic species, but the magnitude varies across traits.

Aquatic paedomorphs and terrestrial direct developers exhibit greater ecomorphological divergence between their limb bone shapes than multiphasic species in those same environments. This likely reflects the narrower range of selective pressures associated with consistent habitat use over ontogeny, whereas multiphasic species are exposed to contrasting physical demands associated with living in water and on land across life stages (Wilson 2005; Bonett and Blair 2017). It also suggests that limb development is not entirely decoupled by metamorphosis, supported by observations of limited anatomical changes between life stages (Ashley-Ross 1992). Therefore, multiphasic species have evolved limb bones influenced by both aquatic and terrestrial demands. Interspecific variation in the timing and duration of life history characteristics that affect time spent in water or on land (i.e., the aquatic larval stage, terrestrial eft stage, or aquatic breeding season) likely influence the relative impacts of the environment and primary agents of selection acting on bone morphology (Weaver et al. 2020; Bonett and Ledbetter 2022; Bonett et al. 2022). For example, many newts have exceptionally stiff bones, possibly reflecting their terrestrial juvenile phases that can last seven years and are absent in other semi-aquatic taxa (Forester and Lykens 1991).

The relative differences in shape, allometry, and evolutionary rates between multiphasic species living in different environments do not always parallel the patterns observed among monophasic species. Many multiphasic species have robust external limb bone shapes not observed in monophasic species and cross-sectional traits that rival direct developers with the most extreme stiffness and density values. These results indicate that multiphasic species are not intermediate versions of monophasic species but instead have their own distinct evolutionary trajectories. Similarly, semi-aquatic lineages are not simply intermediates between aquatic and terrestrial species. These findings suggest that the repeated losses of multiphasic life cycles are unlikely to represent mechanisms for escaping the constraints imposed by complex life cycles. Instead, despite potential life cycle constraints, multiphasic species have highly evolvable limb bones that may have played a role in the reacquisition of an aquatic larval stage and semi-aquatic habitat preferences (Chippindale et al. 2004; Bonett et al. 2014).

### Decoupling of locomotor structures

Weak evolutionary covariance between the external and internal limb bone morphologies are reflected in the independent correlations with habitat and life cycles. We posit that their decoupling has expanded the locomotor capabilities of direct developers and plethodontid salamanders more broadly and, in part, enabled their ecological diversification. Specifically, we propose that the long and gracile plethodontid limb bones increase mobility and locomotor speed (Young et al. 2014), while the internal morphologies can reflect diverse mechanical demands. That is evidenced by the fact that plethodontids occupy diverse microhabitats (i.e., burrows, trees, rocks, caves, streams) (Blankers et al. 2012; Baken and Adams 2019) and employ terrestrial locomotor modes rarely observed outside of plethodontids (e.g., jumping and climbing) (Brown and Deban 2020; Aretz et al. 2022; Huie et al. 2025). In contrast, aquatic paedomorphs have conserved cross-sectional shapes but diverse external shapes, perhaps as the result of relaxed selection on walking kinematics, body shape, and body size in aquatic environments (Ashley-Ross and Bechtel 2004; Azizi and Horton 2004; Bonett and Blair 2017). The only shared external feature of paedomorph limb bones is the reduction or absence of ossified epiphyses, which are formed by cartilage in aquatic salamanders (Molnar 2021b).

Ecological transitions have also influenced the evolutionary correlation between limbs. All ecotypes exhibit relatively strong integration between the external and internal shapes of their limbs (r-pls = 0.88-0.98), possibly reflecting their relatively conserved form of limb-based locomotion (Pierce et al. 2020). Surprisingly, paedomorphs have the strongest integration of external shapes but the weakest covariance between their cross-sectional traits. The latter is aligned with the reduced integration between forelimb and hindlimb lengths among aquatic salamanders and the loss of hindlimbs in aquatic sirens (Ledbetter and Bonett 2019), perhaps reflecting more decoupled mechanical demands. In contrast, direct developers exhibit strong patterns of integration between all humeral and femoral traits, which may reflect the shared load-bearing demands imposed onto the forelimbs and hindlimbs associated with living on land (Kawano et al. 2015; Kawano and Blob 2022).

Multiphasic species also exhibit considerable variation in the evolutionary integration of their forelimbs and hindlimbs. Semi-aquatic species exhibit some of the weakest integration between their external shapes but strongest integration between cross-sectional traits. Because terrestrial multiphasic species do not exhibit these patterns, they are likely dissociated from the demands of terrestrial locomotion and instead reflect unexamined factors. For example, variation in the developmental timing of the forelimbs and hindlimbs may influence patterns of integration in adults. Pond-dwelling larvae exhibit a pronounced lag time between forelimb and hindlimb development, while steam-dwelling larvae (and direct developers) experience less of a lag likely due to an increased need for station holding and underwater walking in streams compared to ponds (Shubin and Wake 2003). Investigations into whether larval development influences levels of integration and if that varies across life stages could yield insights into the factors that generate limb diversity.

### Future directions

In this study we found that interactions between habitat and complex life cycle shape the evolutionary morphology of salamander limb bones. Functional trade-offs between stiffness and density have seemingly promoted stiff yet lightweight bones on land and restricted how these traits covary. Yet, because terrestrial salamanders are relatively small animals, additional work is needed to empirically test whether the forces acting on the limb bones of terrestrial species place demands on the musculoskeletal system strong enough to constrain morphological evolution in the limbs. Walking alone is unlikely to cause bone breakage due to the high safety factors (‘margins against failure’) found in salamanders (Kawano et al. 2013), but other terrestrial activities like running, jumping, or combat may promote the stiffer shapes that we observed (Biewener and Patek 2018).

Additionally, we suggest that complex life cycles promote distinct limb bone morphologies to manage the contrasting aquatic and terrestrial demands experienced within a single lifetime. Future work should examine limb bones over ontogeny to assess the magnitude of change that occurs in morphology, integration, and mechanical properties across metamorphosis and habitat transitions. Doing so will directly address whether metamorphosis can decouple limb bone development between life stages and if complex life cycles necessitate alternative morphologies.

Finally, examining specific clades in greater detail will reveal more nuanced factors affecting limb evolution. For instance, it is unclear to what extent sexual dimorphism and courtship behaviors in salamandrids influence the properties of their limb bones (Reinhard et al. 2015). We also recognize that the semi-aquatic ecotype as defined in this study may be overly broad because it includes species that spend most of their lives at the water’s edge as well as newts with three life stages and multiple transitions between aquatic and terrestrial stages. We suspect that the different semi-aquatic strategies have distinct consequences on limb bone evolution, and encourage future studies to test this hypothesis at a finer scale than what was focused on in this study. In sum, further investigation into how the biomechanical properties of salamander limb bones relate to life cycle strategy, locomotion, and other functional applications would provide a deeper understanding of the selective pressures governing musculoskeletal evolution.

## ACKNOWLEDGEMENTS

We thank P. Millers for providing access to Burke Museum specimens amidst the COVID-19 pandemic, D. Grossnickle for retrieving the specimens, and A. Summers and the Karel F. Liem Bio-Imaging facility for free access to a micro-CT scanner. We also thank H. Martens, R. Nagesan, and G. Schneider for providing CT scans of UMMZ specimens. We thank D. Caetano for helping with the *ratematrix* analyses. We also thank two anonymous reviewers for their helpful feedback on earlier drafts of this manuscript.

## AUTHOR CONTRIBUTIONS

J.M.H. and S.M.K. designed research; J.M.H. performed research; J.M.H. and R.A.P. analyzed data; and J.M.H., R.A.P., and S.M.K. wrote the paper.

## DATA AVAILABILITY

The morphological data and R script files used for this study can be found on GitHub (https://github.com/jmhuie/Salamander_Limb_Bone_Evo).

