## Supplemental Information for "Habitat and complex life cycles promote morphological diversity in salamander limb bones"

**In this document:**

*Supplemental Figures*

- Figure S1: The template bones with the landmarking scheme.
- Figure S2: Allometric variation in limb bone morphology in relationship to body size.

*Supplemental Tables*

- Table S1: List of 133 species sampled in this study and specimen metadata.
- Table S2: Morphological disparity by ecotype.
- Table S3: Phylogenetic ANOVAs results.
- Table S4: Pairwise comparisons of limb bone morphologies between ecotypes.
- Table S5: Phylogenetic paired t-tests comparing forelimbs and hindlimbs.
- Table S6: Evolutionary rates of external limb shape by ecotype.
- Table S7: Fit of the 26 *hOUwie* models for humeral stiffness with sirens.
- Table S8: Fit of the 26 *hOUwie* models for humeral stiffness without sirens.
- Table S9: Fit of the 26 *hOUwie* models for femoral stiffness.
- Table S10: Fit of the 26 *hOUwie* models for humeral density with sirens.
- Table S11: Fit of the 26 *hOUwie* models for humeral density without sirens.
- Table S12: Fit of the 26 *hOUwie* models for femoral density.
- Table S13: Model averaged evolutionary rates and cross-sectional trait optima.

*Supplemental References*

**Not in this document:**

The morphological data and R script files used for this study can be found on GitHub (<https://github.com/jmhuie/Salamander_Limb_Bone_Evo>).

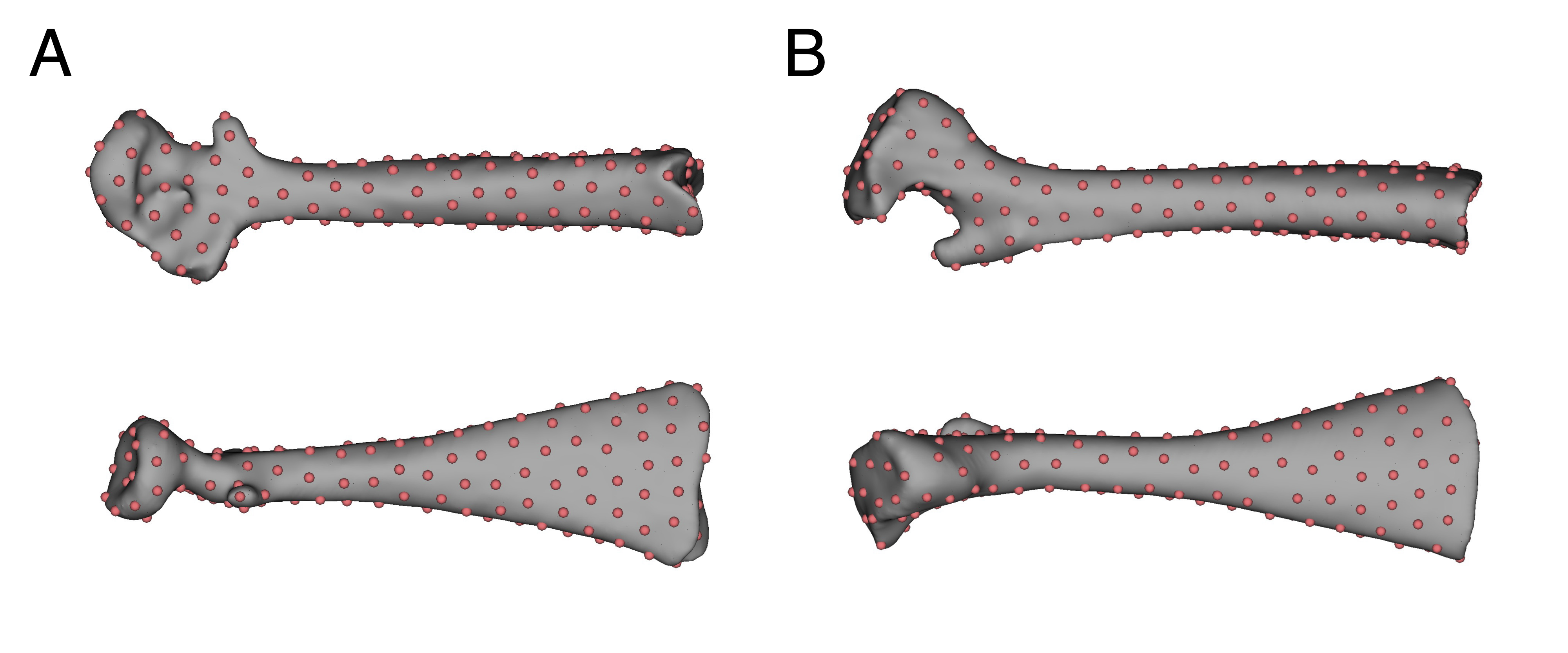

Figure S1. The template bones with the landmarking scheme. Dorsal and lateral views of A) the humerus of *Aneides hardii* with 193 pseudolandmarks and B) the femur of *Plethodon elongatus* with 190 pseudolandmarks. The templates were selected based on which species were closest to the mean shape in preliminary analyses. The number of pseudolandmarks were selected by visually evaluating the number of points needed to comprehensively sample the surface of the most extreme bones. We then performed sensitivity analyses to assess whether our results were robust to the number of landmarks using the “LaSEC” (Landmark Sampling Evaluation Curve) function in the LaMBDA R package (1). This method iteratively subsamples the number of landmarks in the data set and assesses how well the different subsamples converge on similar patterns of morphological variation. We found that for both the humeral and femoral datasets that ~100 landmarks (and more) effectively captured the same patterns of variation as the original ~190 landmark datasets. That was indicated by plateaus in model fit and suggests that our landmarks datasets are sufficient for capturing the primary axes of shape variation.

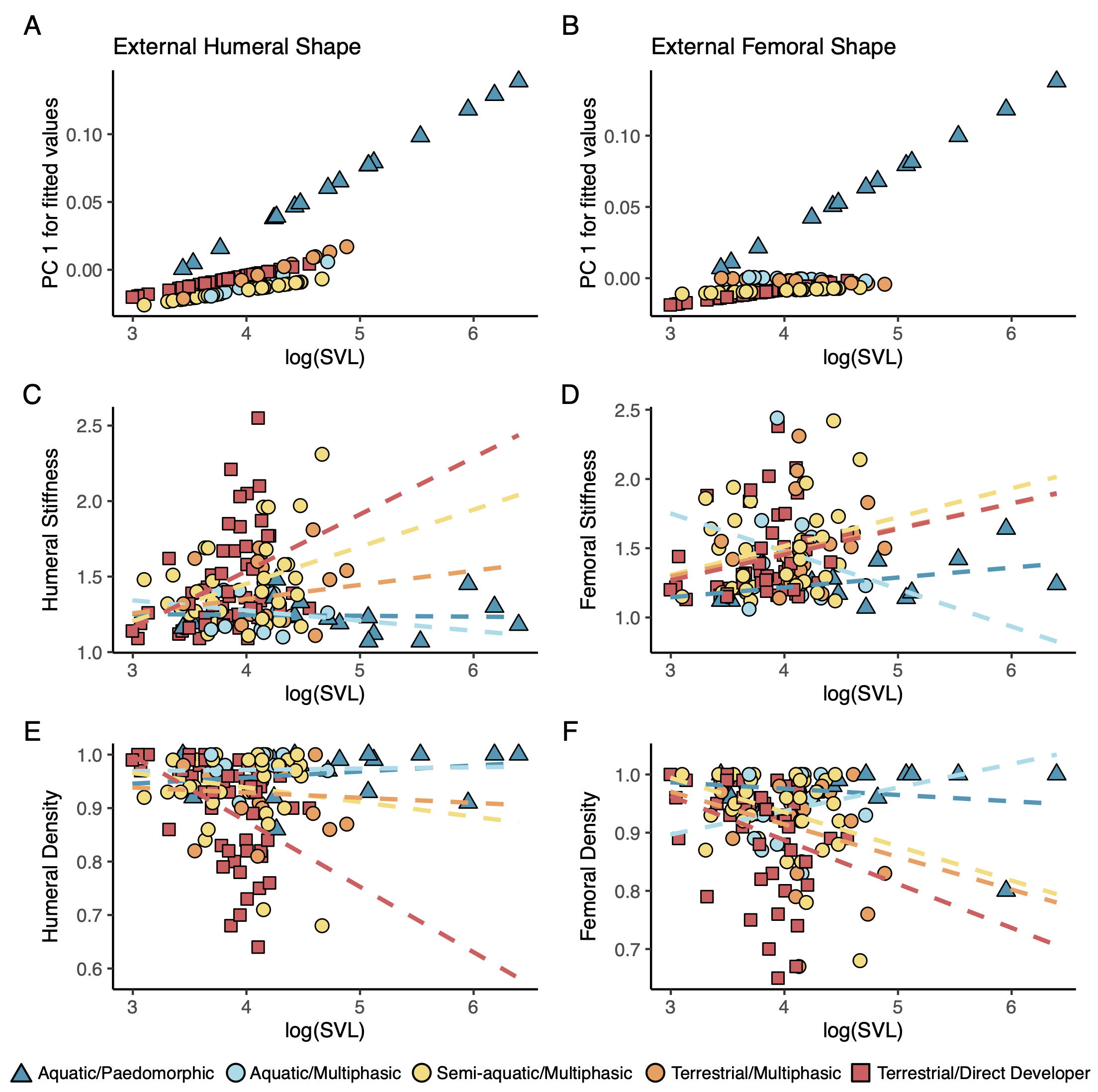

Figure S2. Allometric variation in external limb bone shape and cross-sectional traits in relationship to body size. External shape plots (A, B) were made using the “plotAllometry” function in the *geomorph* R package with the “PredLine” argument. This method takes the fitted values from regressions previously performed with “procD.pgls” and plots the first principal component of the fitted values against body size. The cross-sectional trait plots depict the ordinary least squares regression lines between the trait values and body size, as estimated with the “geom_smooth” function for *ggplot2*. Note that phylogenetic ANOVAs indicated the ecotype did not have a significant effect on the allometric relationships of the cross-sectional traits from the femora.

**Table S1.** List of 133 species sampled in this study with their specimen information, link the scans on a digital repository, and habitat and life cycle classifications. Some multiphasic species exhibit facultative paedomorphism, in which cases habitat and life cycle classifications match the ecology of the specimen sampled. * = terrestrial adult sampled (for facultatively paedomorphic species), + = paedomorphic adult sampled (for facultatively paedomorphic species); A = Aquatic, SAq = Semi-aquatic, and T = Terrestrial; pd = Paedomorphic, bi = multiphasic, and dd = Direct developer.

| **Species** | **Specimen Number** | **Family** | **Data Repository** | **Habitat** | **Life Cycle** | **Ref** |
| --- | --- | --- | --- | --- | --- | --- |
| *Ambystoma dumerilii* | UF 43549 | Ambystomatidae | <https://www.morphosource.org/media/000140391> | Aq | pd | (2) |
| *Ambystoma mabeei* | UWBM 4987 | Ambystomatidae | <https://www.morphosource.org/media/000629427> | T | bi | (3) |
| *Ambystoma maculatum* | UF 26607 | Ambystomatidae | <https://www.morphosource.org/media/000141549> | T | bi | (2) |
| *Ambystoma mavortium* | UF 158300 | Ambystomatidae | <https://www.morphosource.org/media/000038426> | T | bi | (2) |
| *Ambystoma mexicanum* | UWBM 7027 | Ambystomatidae | <https://www.morphosource.org/media/000629432> | Aq | pd | (2) |
| *Ambystoma ordinarium+* | UCM 22117 | Ambystomatidae | <https://www.morphosource.org/media/000438897> | Aq | pd | (2) |
| *Ambystoma rosaceum+* | UCM 66358 | Ambystomatidae | <https://www.morphosource.org/media/000387225> | Aq | pd | (2) |
| *Ambystoma talpoideum** | UWBM 4989 | Ambystomatidae | <https://www.morphosource.org/media/000629437> | T | bi | (2) |
| *Ambystoma tigrinum** | UMZC R.506 | Ambystomatidae | <https://www.morphosource.org.media/000515776> | T | bi | (2) |
| *Amphiuma tridactylum* | NHMUK II.II.2.1a | Amphiumidae | [https://www.morphosource.org/media/000080420](about:blank) | Aq | pd | (2) |
| *Andrias japonicus* | FMNH 31536 | Cryptobranchidae | <https://www.morphosource.org/media/000077893> | Aq | pd | (2) |
| *Aneides aeneus* | MVZ 52960 | Plethodontidae | <https://www.morphosource.org/media/000165921> | T | dd | (2) |
| *Aneides ferreus* | MVZ 145629 | Plethodontidae | <https://www.morphosource.org/media/000165931> | T | dd | (2) |
| *Aneides flavipunctatus* | MVZ 123824 | Plethodontidae | <https://www.morphosource.org/media/000165897> | T | dd | (2) |
| *Aneides hardii* | MVZ 220872 | Plethodontidae | <https://www.morphosource.org/media/000165936> | T | dd | (2) |
| *Aneides lugubris* | MVZ 249828 | Plethodontidae | <https://www.morphosource.org/media/000049486> | T | dd | (2) |
| *Aneides vagrans* | MVZ 242271 | Plethodontidae | <https://www.morphosource.org/media/000165925> | T | dd | (2) |
| *Aquiloeurycea cephalica* | UCM 8183 | Plethodontidae | <https://www.morphosource.org/media/000386875> | T | dd | (2) |
| *Aquiloeurycea galeanae* | UCM 47446 | Plethodontidae | <https://www.morphosource.org/media/000453564> | T | dd | (2) |
| *Batrachoseps wrighti* | UWBM 4624 | Plethodontidae | <https://www.morphosource.org/media/000629451> | T | dd | (2) |
| *Batrachuperus pinchonii* | MVZ 234909 | Hynobiidae | <https://www.morphosource.org/media/000049473> | Aq | bi | (2) |
| *Bolitoglossa diminuta* | MVZ 207052 | Plethodontidae | <https://www.morphosource.org/media/000480968> | T | dd | (2) |
| *Bolitoglossa dofleini* | UMMZ 89119 | Plethodontidae | <https://doi.org/10.7302/4m0x-5x26> | T | dd | (2) |
| *Bolitoglossa franklini* | UF 172966 | Plethodontidae | <https://www.morphosource.org/media/000036165> | T | dd | (2) |
| *Bolitoglossa gracilis* | MVZ 229170 | Plethodontidae | <https://www.morphosource.org/media/000480977> | T | dd | (2) |
| *Bolitoglossa hartwegi* | MVZ 131711 | Plethodontidae | <https://www.morphosource.org/media/000480956> | T | dd | (2) |
| *Bolitoglossa heiroreias* | YPM 7242 | Plethodontidae | <https://www.morphosource.org/media/000494816> | T | dd | (2) |
| *Bolitoglossa huehuetenanguensis* | MVZ 269875 | Plethodontidae | <https://www.morphosource.org/media/000477973> | T | dd | (2) |
| *Bolitoglossa mulleri* | UCM 34847 | Plethodontidae | <https://www.morphosource.org/media/000386985> | T | dd | (2) |
| *Bolitoglossa porrasorum* | UF 156522 | Plethodontidae | <https://www.morphosource.org/media/000049711> | T | dd | (2) |
| *Bolitoglossa robinsoni* | MVZ 222477 | Plethodontidae | <https://www.morphosource.org/media/000476232> | T | dd | (2) |
| *Bolitoglossa rufescens* | UCM 22139 | Plethodontidae | <https://www.morphosource.org/media/000386937> | T | dd | (2) |
| *Bradytriton silus* | MVZ 265367 | Plethodontidae | <https://www.morphosource.org/media/000049491> | T | dd | (2) |
| *Calotriton asper** | MCZ 107494 | Salamandridae | <https://www.morphosource.org/media/000043538> | Aq | bi | (2) |
| *Chioglossa lusitanica* | MVZ 71635 | Salamandridae | <https://www.morphosource.org/media/000055891> | SAq | bi | (2) |
| *Chiropterotriton chiropterus* | UMMZ 115115a | Plethodontidae | <https://doi.org/10.7302/amrc-pj88> | T | dd | (2) |
| *Chiropterotriton magnipes* | MVZ 128260 | Plethodontidae | <https://www.morphosource.org/media/000492429> | T | dd | (2) |
| *Cryptobranchus alleganiensis* | UF 88726 | Cryptobranchidae | <https://www.morphosource.org/media/000036426> | Aq | pd | (2) |
| *Cynops pyrrhogaster* | UF 92871 | Salamandridae | <https://www.morphosource.org/media/000038137> | Aq | bi | (2) |
| *Dendrotriton megarhinus* | MVZ 206209 | Plethodontidae | <https://www.morphosource.org/media/000476226> | T | dd | (2) |
| *Desmognathus amphileucus* | BMNH 2021.7551 | Plethodontidae | <https://www.morphosource.org/media/000630363> | SAq | bi | (2) |
| *Desmognathus aureatus* | AMNH A-194384 | Plethodontidae | <https://www.morphosource.org/media/000630330> | Aq | bi | (2) |
| *Desmognathus fuscus* | OUMNH 10037 | Plethodontidae | <https://www.morphosource.org/media/000163357> | SAq | bi | (2) |
| *Desmognathus monticola* | AMNH A-195565 | Plethodontidae | <https://www.morphosource.org/media/000561226> | SAq | bi | (2) |
| *Desmognathus wrighti* | AMNH A-196154 | Plethodontidae | <https://www.morphosource.org/media/000630364> | T | dd | (2) |
| *Dicamptodon copei+* | UWBM 74 | Dicamptodontidae | <https://www.morphosource.org/media/000629444> | Aq | pd | (2) |
| *Dicamptodon ensatus** | UF 149094 | Dicamptodontidae | <https://www.morphosource.org/media/000133974> | T | bi | (2) |
| *Dicamptodon tenebrosus** | UWBM 1808 | Dicamptodontidae | [https://www.morphosource.org//media/000630158](https://www.morphosource.org/media/000630158) | T | bi | (2) |
| *Echinotriton andersoni* | UMMZ 174420 | Salamandridae | <https://www.morphosource.org/media/000057178> | T | bi | (2) |
| *Ensatina eschscholtzii* | MVZ 237515 | Plethodontidae | <https://www.morphosource.org/media/000049478> | T | dd | (2) |
| *Euproctus platycephalus** | MVZ 129572 | Salamandridae | <https://www.morphosource.org/media/000049445> | Aq | bi | (2) |
| *Eurycea arenicola* | NCSM 24613 | Plethodontidae | <https://www.morphosource.org/media/000039696> | SAq | bi | (3) |
| *Eurycea longicauda* | UWBM 4582 | Plethodontidae | <https://www.morphosource.org/media/000630159> | T | bi | (2) |
| *Eurycea lucifuga* | KU 51584 | Plethodontidae | <https://www.morphosource.org/media/000059603> | T | bi | (2) |
| *Eurycea neotenes* | NHMUK 1957.1.7.88-89 | Plethodontidae | <https://www.morphosource.org/media/000517449> | Aq | pd | (2) |
| *Eurycea wallacei* | YPM 13629 | Plethodontidae | <https://www.morphosource.org/media/000050172> | Aq | pd | (2) |
| *Eurycea wilderae* | GMNH 43944 | Plethodontidae | <https://www.morphosource.org/media/000607347> | SAq | bi | (2) |
| *Gyrinophilus porphyriticus* | UF 64645 | Plethodontidae | <https://www.morphosource.org/media/000036213> | SAq | bi | (2) |
| *Hemidactylium scutatum* | UMMZ 246927 | Plethodontidae | <https://doi.org/10.7302/c2j1-sy06> | T | bi | (2) |
| *Hydromantes genei* | MVZ 205030 | Plethodontidae | <https://www.morphosource.org/media/000049455> | T | dd | (2) |
| *Hydromantes platycephalus* | MVZ 236798 | Plethodontidae | <https://www.morphosource.org/media/000049475> | T | dd | (2) |
| *Hydromantes samweli* | MVZ 170637 | Plethodontidae | <https://www.morphosource.org/media/000074070> | T | dd | (2) |
| *Hynobius naevius* | OUMNH 6990 | Hynobiidae | <https://www.morphosource.org/media/000099645> | T | bi | (4) |
| *Hynobius nebulosus* | UF 24315 | Hynobiidae | <https://www.morphosource.org/media/000133873> | T | bi | (2) |
| *Hypselotriton orientalis* | MCZ 151391 | Salamandridae | <https://www.morphosource.org/media/000166445> | Aq | bi | (2) |
| *Hypselotriton wolterstorffi** | MCZ 8156 | Salamandridae | <https://www.morphosource.org/media/000100232> | Aq | bi | (2) |
| *Ichthyosaura alpestris** | UF 13359 | Salamandridae | <https://www.morphosource.org/media/000037673> | SAq | bi | (2) |
| *Isthmura bellii* | UMMZ 143736 | Plethodontidae | <https://doi.org/10.7302/x6qs-9372> | T | dd | (2) |
| *Ixalotriton parvus* | MVZ 177824 | Plethodontidae | <https://www.morphosource.org/media/000491034> | T | dd | (2) |
| *Laotriton laoensis* | NCSM 79785 | Salamandridae | <https://www.morphosource.org/media/000039516> | SAq | bi | (2) |
| *Lissotriton boscai* | MCZ 125997 | Salamandridae | <https://www.morphosource.org/media/000471946> | SAq | bi | (2) |
| *Lissotriton helveticus** | UF 38991 | Salamandridae | <https://www.morphosource.org/media/000036187> | SAq | bi | (2) |
| *Lissotriton italicus** | MCZ 7367 | Salamandridae | <https://www.morphosource.org/media/000470347> | SAq | bi | (2) |
| *Lissotriton montandoni* | MCZ 107643 | Salamandridae | <https://www.morphosource.org/media/000468776> | SAq | bi | (2) |
| *Lissotriton vulgaris** | UF 39025 | Salamandridae | <https://www.morphosource.org/media/000036416> | SAq | bi | (2) |
| *Liua shihi* | MVZ 231146 | Hynobiidae | <https://www.morphosource.org/media/000049467> | Aq | bi | (2) |
| *Mertensiella caucasica* | MVZ 218721 | Salamandridae | <https://www.morphosource.org/media/000049456> | T | bi | (2) |
| *Necturus lewisi* | NCSM 19905 | Proteidae | <https://www.morphosource.org/media/000039520> | Aq | pd | (2) |
| *Necturus maculosus* | UMZC R.16130 | Proteidae | <https://www.morphosource.org/media/000163870> | Aq | pd | (2) |
| *Neurergus crocatus* | MVZ 236763 | Salamandridae | <https://www.morphosource.org/media/000059420> | SAq | bi | (2) |
| *Notophthalmus viridescens** | UF 84517 | Salamandridae | <https://www.morphosource.org/media/000036288> | SAq | bi | (2) |
| *Nototriton mime* | MVZ 269306 | Plethodontidae | <https://www.morphosource.org/media/000492465> | T | dd | (2) |
| *Nototriton picadoi* | UMMZ 238369 | Plethodontidae | <https://doi.org/10.7302/6dqg-pn61> | T | dd | (2) |
| *Nyctanolis pernix* | MVZ 263972 | Plethodontidae | <https://www.morphosource.org/media/000049488> | T | dd | (2) |
| *Oedipina maritima* | MVZ 219997 | Plethodontidae | <https://www.morphosource.org/media/000491046> | T | dd | (2) |
| *Oedipina savagei* | MVZ 229360 | Plethodontidae | <https://www.morphosource.org/media/000491052> | T | dd | (2) |
| *Oedipina taylori* | MVZ 267200 | Plethodontidae | <https://www.morphosource.org/media/000480989> | T | dd | (2) |
| *Ommatotriton vittatus** | MVZ 219525 | Salamandridae | <https://www.morphosource.org/media/000049458> | SAq | bi | (2) |
| *Onychodactylus japonicus* | CAS 26711 | Hynobiidae | <https://www.morphosource.org/media/000050024> | SAq | bi | (3) |
| *Pachyhynobius shangchengensis* | CAS 194247 | Hynobiidae | <https://www.morphosource.org/media/000059250> | Aq | bi | (2) |
| *Pachytriton brevipes* | MCZ 22346 | Salamandridae | <https://www.morphosource.org/media/000167778> | Aq | bi | (2) |
| *Paradactylodon persicus* | MVZ 241494 | Hynobiidae | <https://www.morphosource.org/media/000049481> | SAq | bi | (3) |
| *Paramesotriton deloustali* | FMNH 259125 | Salamandridae | <https://www.morphosource.org/media/000048649> | SAq | bi | (3) |
| *Paramesotriton labiatus* | UF 157219 | Salamandridae | <https://www.morphosource.org/media/000036675> | SAq | bi | (3) |
| *Parvimolge townsendi* | UMMZ 118145 | Plethodontidae | <https://doi.org/10.7302/88rt-vz27> | T | dd | (2) |
| *Plethodon cinereus* | UF 43538 | Plethodontidae | <https://www.morphosource.org/media/000100264> | T | dd | (2) |
| *Plethodon dunni* | MVZ 220000 | Plethodontidae | <https://www.morphosource.org/media/000165917> | T | dd | (2) |
| *Plethodon elongatus* | MVZ 84856 | Plethodontidae | <https://www.morphosource.org/media/000165927> | T | dd | (2) |
| *Plethodon glutinosus* | OUMNH 10039 | Plethodontidae | <https://www.morphosource.org/media/000099728> | T | dd | (2) |
| *Plethodon grobmani* | UF 16267 | Plethodontidae | <https://www.morphosource.org/media/000049700> | T | dd | (2) |
| *Plethodon montanus* | UWBM 4985 | Plethodontidae | <https://www.morphosource.org/media/000630122> | T | dd | (2) |
| *Plethodon petraeus* | USNM 267168 | Plethodontidae | <https://www.morphosource.org/media/000725038> | T | dd | (2) |
| *Plethodon vehiculum* | UWBM 693 | Plethodontidae | <https://www.morphosource.org/media/000561214> | T | dd | (2) |
| *Pleurodeles waltl** | CAS 138786 | Salamandridae | <https://www.morphosource.org/media/000050010> | Aq | bi | (2) |
| *Proteus anguinus* | MVZ 244076 | Proteidae | <https://www.morphosource.org/media/000049506> | Aq | pd | (2) |
| *Pseudobranchus axanthus* | UF 167203 | Sirenidae | <https://www.morphosource.org/media/000049081> | Aq | pd | (2) |
| *Pseudobranchus striatus* | UF 179649 | Sirenidae | <https://www.morphosource.org/media/000049084> | Aq | pd | (2) |
| *Pseudoeurycea cochranae* | UCM 60247 | Plethodontidae | <https://www.morphosource.org/media/000413572> | T | dd | (2) |
| *Pseudoeurycea leprosa* | MVZ 158765 | Plethodontidae | <https://www.morphosource.org/media/000049448> | T | dd | (2) |
| *Pseudoeurycea melanomolga* | UCM 57165 | Plethodontidae | <https://www.morphosource.org/media/000387656> | T | dd | (2) |
| *Pseudohynobius flavomaculatus* | MVZ 231152 | Hynobiidae | <https://www.morphosource.org/media/000049469> | T | bi | (2) |
| *Pseudotriton ruber* | UMMZ 194448 | Plethodontidae | <https://doi.org/10.7302/23qf-rd17> | SAq | bi | (2) |
| *Ranodon sibiricus* | UMMZ 127463 | Hynobiidae | <https://www.morphosource.org/media/000057171> | SAq | bi | (3, 4) |
| *Rhyacotriton kezeri* | UWBM 3147 | Rhyacotritonidae | <https://www.morphosource.org/media/000630319> | SAq | bi | (2) |
| *Rhyacotriton olympicus* | UMMZ 135501 | Rhyacotritonidae | <https://www.morphosource.org/media/000057174> | SAq | bi | (2) |
| *Rhyacotriton variegatus* | UWBM 6082 | Rhyacotritonidae | <https://www.morphosource.org/media/000630324> | SAq | bi | (3) |
| *Salamandra algira* | NHMUK 1889.12.7.6-7 | Salamandridae | <https://www.morphosource.org/media/000163613> | T | bi | (2) |
| *Salamandra salamandra* | MCZ 2796 | Salamandridae | <https://www.morphosource.org/media/000165871> | T | bi | (2) |
| *Salamandrella keyserlingii* | MVZ 222337 | Hynobiidae | <https://www.morphosource.org/media/000049460> | T | bi | (2) |
| *Salamandrina terdigitata* | MVZ 178849 | Salamandridae | <https://www.morphosource.org/media/000055881> | T | bi | (2) |
| *Siren intermedia* | UF 123993 | Sirenidae | <https://www.morphosource.org/media/000025580> | Aq | pd | (2) |
| *Siren lacertina* | UMZC R.763 | Sirenidae | <https://www.morphosource.org/media/000163656> | Aq | pd | (2) |
| *Stereochilus marginatus* | UMMZ 126557a | Plethodontidae | <https://doi.org/10.7302/14qx-zd60> | Aq | bi | (2) |
| *Taricha granulosa** | MCZ 150332 | Salamandridae | <https://www.morphosource.org/media/000166453> | SAq | bi | (3) |
| *Taricha rivularis* | MCZ 22496 | Salamandridae | <https://www.morphosource.org/media/000166373> | SAq | bi | (3) |
| *Taricha torosa* | UMZC R.16085 | Salamandridae | <https://www.morphosource.org/media/000515788> | SAq | bi | (2) |
| *Thorius macdougalli* | UMMZ 119705 | Plethodontidae | <https://doi.org/10.7302/ejfc-fy07> | T | dd | (2) |
| *Triturus carnifex** | MCZ 126041 | Salamandridae | <https://www.morphosource.org/media/000469560> | SAq | bi | (2) |
| *Triturus cristatus** | FMNH 84926 | Salamandridae | <https://www.morphosource.org/media/000165162> | SAq | bi | (2) |
| *Tylototriton panhai* | NCSM 82961 | Salamandridae | <https://www.morphosource.org/media/000039521> | SAq | bi | (5) |
| *Tylototriton taliangensis* | CAS 195131 | Salamandridae | <https://www.morphosource.org/media/000059249> | SAq | bi | (3) |
| *Tylototriton verrucosus* | CAS 242371 | Salamandridae | <https://www.morphosource.org/media/000050023> | SAq | bi | (2) |
| *Urspelerpes brucei* | Dave Beamer 6337 | Plethodontidae | <https://www.morphosource.org/media/000120675> | SAq | bi | (3) |

**Table S2.** Morphological disparity by ecotype. All humeral analyses were performed with and without the sirens included in the dataset. A = Aquatic, SAq = Semi-aquatic, and T = Terrestrial; pd = Paedomorphic, bi = multiphasic, and dd = Direct developer.

|  | **Aq\|pd** | **Aq\|bi** | **SAq\|bi** | **T\|bi** | **T\|dd** |
| --- | --- | --- | --- | --- | --- |
| External Humerus Shape (with sirens) | 0.0111 | 0.0053 | 0.0066 | 0.0072 | 0.0040 |
| External Humerus Shape (no sirens) | 0.0090 | 0.0053 | 0.0066 | 0.0072 | 0.0040 |
| External Femur Shape | 0.0088 | 0.0043 | 0.0052 | 0.0059 | 0.0047 |
| Humeral Stiffness (with sirens) | 0.0127 | 0.0164 | 0.0761 | 0.0423 | 0.1067 |
| Humeral Stiffness (no sirens) | 0.0083 | 0.0164 | 0.0761 | 0.0423 | 0.1067 |
| Femoral Stiffness | 0.0256 | 0.1317 | 0.1022 | 0.1004 | 0.0811 |
| Humeral Density (with sirens) | 0.0017 | 0.0012 | 0.0055 | 0.0032 | 0.0089 |
| Humeral Density (no sirens) | 0.0010 | 0.0012 | 0.0055 | 0.0032 | 0.0089 |
| Femoral Density | 0.0028 | 0.0034 | 0.0049 | 0.0076 | 0.0081 |

**Table S3.** Results of the phylogenetic ANOVAs performed on the external and internal limb bone traits. Each analysis was performed with the format: trait ~ log(SVL)*Ecotype. All humeral analyses were performed with and without the sirens included in the dataset. Bold values indicate statistical significance (p < 0.05).

|  | **Covariate** | **Df** | **SS** | **MS** | **R^2^** | **F** | **Z** | **p-value** |
| --- | --- | --- | --- | --- | --- | --- | --- | --- |
| External Humerus Shape (with sirens) | log(SVL) | 1 | 0.001 | 0.001 | 0.031 | 4.519 | 4.286 | **0.001** |
|  | Ecotype | 4 | 0.002 | 0.001 | 0.079 | 2.891 | 4.218 | **0.001** |
|  | log(SVL)*Ecotype | 4 | 0.001 | 0.000 | 0.047 | 1.711 | 2.672 | **0.002** |
| External Humerus Shape (no sirens) | log(SVL) | 1 | 0.001 | 0.001 | 0.036 | 5.276 | 4.836 | **0.001** |
|  | Ecotype | 4 | 0.002 | 0.001 | 0.083 | 3.028 | 4.573 | **0.001** |
|  | log(SVL)*Ecotype | 4 | 0.002 | 0.000 | 0.060 | 2.184 | 3.632 | **0.001** |
| External Femur Shape | log(SVL) | 1 | 0.001 | 0.001 | 0.027 | 3.841 | 3.404 | **0.002** |
|  | Ecotype | 4 | 0.002 | 0.001 | 0.090 | 3.251 | 4.751 | **0.001** |
|  | log(SVL)*Ecotype | 4 | 0.001 | 0.000 | 0.056 | 2.002 | 3.112 | **0.001** |
| Humeral Stiffness  (with sirens) | log(SVL) | 1 | 0.010 | 0.010 | 0.001 | 0.158 | -0.486 | 0.680 |
|  | Ecotype | 4 | 1.405 | 0.351 | 0.139 | 5.376 | 3.063 | **0.003** |
|  | log(SVL)*Ecotype | 4 | 0.688 | 0.172 | 0.068 | 2.631 | 1.739 | **0.040** |
| Humeral Stiffness  (no sirens) | log(SVL) | 1 | 0.042 | 0.042 | 0.004 | 0.630 | 0.268 | 0.427 |
|  | Ecotype | 4 | 1.451 | 0.363 | 0.145 | 5.444 | 3.293 | **0.001** |
|  | log(SVL)*Ecotype | 4 | 0.592 | 0.148 | 0.059 | 2.220 | 1.432 | 0.075 |
| Femoral Stiffness | log(SVL) | 1 | 0.087 | 0.087 | 0.007 | 0.955 | 0.494 | 0.331 |
|  | Ecotype | 4 | 0.909 | 0.227 | 0.075 | 2.499 | 1.626 | 0.053 |
|  | log(SVL)*Ecotype | 4 | 0.243 | 0.061 | 0.020 | 0.669 | -0.276 | 0.606 |
| Humeral Density  (with sirens) | log(SVL) | 1 | 0.000 | 0.000 | 0.000 | 0.011 | -1.414 | 0.897 |
|  | Ecotype | 4 | 0.121 | 0.030 | 0.144 | 5.723 | 3.248 | **0.001** |
|  | log(SVL)*Ecotype | 4 | 0.072 | 0.018 | 0.085 | 3.402 | 2.168 | **0.013** |
| Humeral Density  (no sirens) | log(SVL) | 1 | 0.001 | 0.001 | 0.001 | 0.151 | -0.506 | 0.691 |
|  | Ecotype | 4 | 0.132 | 0.033 | 0.160 | 6.195 | 3.528 | **0.001** |
|  | log(SVL)*Ecotype | 4 | 0.059 | 0.015 | 0.072 | 2.780 | 1.803 | **0.035** |
| Femoral Density | log(SVL) | 1 | 0.005 | 0.005 | 0.006 | 0.862 | 0.444 | 0.362 |
|  | Ecotype | 4 | 0.102 | 0.026 | 0.118 | 4.142 | 2.686 | **0.003** |
|  | log(SVL)*Ecotype | 4 | 0.024 | 0.006 | 0.028 | 0.976 | 0.137 | 0.441 |

**Table S4.** Results of the pairwise comparisons of external and internal limb bone morphologies between ecotypes. Most analyses tested for differences in mean shape between ecotypes, but some tested for differences in allometric slope. Comparisons were only performed for significant covariates indicated in Table S3, except for femoral stiffness where ecotype almost had significant effect (p = 0.053). A = Aquatic, SAq = Semi-aquatic, and T = Terrestrial; pd = Paedomorphic, bi = multiphasic, and dd = Direct developer. Bold values indicate statistical significance (p < 0.05).

|  | **Comparison** | **d** | **UCL (95%)** | **Z** | **p-value** |
| --- | --- | --- | --- | --- | --- |
| Humerus External Shape: Mean (with sirens) | Aq\|pd vs Aq\|bi | 0.061 | 0.052 | 2.521 | **0.005** |
|  | Aq\|pd vs SAq\|bi | 0.061 | 0.046 | 2.937 | **0.001** |
|  | Aq\|pd vs T\|bi | 0.058 | 0.039 | 2.780 | **0.002** |
|  | Aq\|pd vs T\|dd | 0.064 | 0.075 | 0.818 | 0.200 |
|  | Aq\|bi vs SAq\|bi | 0.028 | 0.032 | 0.971 | 0.165 |
|  | Aq\|bi vs T\|bi | 0.031 | 0.045 | -0.507 | 0.687 |
|  | Aq\|bi vs T\|dd | 0.042 | 0.073 | -1.179 | 0.891 |
|  | SAq\|bi vs T\|bi | 0.030 | 0.039 | 0.244 | 0.409 |
|  | SAq\|bi vs T\|dd | 0.038 | 0.071 | -1.360 | 0.913 |
|  | T\|bi vs T\|dd | 0.038 | 0.072 | -1.670 | 0.956 |
| Humerus External Shape: Allometry (with sirens) | Aq\|pd vs Aq\|bi | 0.088 | 0.115 | 0.465 | 0.306 |
|  | Aq\|pd vs SAq\|bi | 0.064 | 0.052 | 2.915 | **0.002** |
|  | Aq\|pd vs T\|bi | 0.097 | 0.067 | 3.099 | **0.001** |
|  | Aq\|pd vs T\|dd | 0.061 | 0.055 | 2.132 | **0.017** |
|  | Aq\|bi vs SAq\|bi | 0.081 | 0.120 | -0.042 | 0.497 |
|  | Aq\|bi vs T\|bi | 0.101 | 0.127 | 0.608 | 0.267 |
|  | Aq\|bi vs T\|dd | 0.082 | 0.123 | -0.178 | 0.566 |
|  | SAq\|bi vs T\|bi | 0.089 | 0.077 | 2.337 | **0.013** |
|  | SAq\|bi vs T\|dd | 0.045 | 0.064 | -0.551 | 0.692 |
|  | T\|bi vs T\|dd | 0.089 | 0.075 | 2.477 | **0.009** |
| Humerus External Shape: Mean (no sirens) | Aq\|pd vs Aq\|bi | 0.061 | 0.052 | 2.494 | **0.006** |
|  | Aq\|pd vs SAq\|bi | 0.060 | 0.046 | 2.865 | **0.001** |
|  | Aq\|pd vs T\|bi | 0.058 | 0.039 | 2.584 | **0.004** |
|  | Aq\|pd vs T\|dd | 0.065 | 0.075 | 1.053 | 0.154 |
|  | Aq\|bi vs SAq\|bi | 0.028 | 0.035 | 0.720 | 0.239 |
|  | Aq\|bi vs T\|bi | 0.032 | 0.046 | -0.461 | 0.683 |
|  | Aq\|bi vs T\|dd | 0.042 | 0.071 | -1.118 | 0.868 |
|  | SAq\|bi vs T\|bi | 0.030 | 0.039 | 0.288 | 0.389 |
|  | SAq\|bi vs T\|dd | 0.038 | 0.067 | -1.104 | 0.857 |
|  | T\|bi vs T\|dd | 0.038 | 0.072 | -1.590 | 0.950 |
| Humerus External Shape: Allometry (no sirens) | Aq\|pd vs Aq\|bi | 0.097 | 0.126 | 0.811 | 0.204 |
|  | Aq\|pd vs SAq\|bi | 0.076 | 0.052 | 3.899 | **0.001** |
|  | Aq\|pd vs T\|bi | 0.103 | 0.067 | 3.418 | **0.001** |
|  | Aq\|pd vs T\|dd | 0.073 | 0.055 | 3.143 | **0.001** |
|  | Aq\|bi vs SAq\|bi | 0.081 | 0.125 | 0.003 | 0.474 |
|  | Aq\|bi vs T\|bi | 0.101 | 0.131 | 0.614 | 0.277 |
|  | Aq\|bi vs T\|dd | 0.081 | 0.127 | -0.165 | 0.565 |
|  | SAq\|bi vs T\|bi | 0.089 | 0.074 | 2.567 | **0.004** |
|  | SAq\|bi vs T\|dd | 0.044 | 0.062 | -0.523 | 0.688 |
|  | T\|bi vs T\|dd | 0.090 | 0.074 | 2.735 | **0.004** |
| Femur External Shape: Mean | Aq\|pd vs Aq\|bi | 0.060 | 0.053 | 2.241 | **0.015** |
|  | Aq\|pd vs SAq\|bi | 0.067 | 0.047 | 2.996 | **0.001** |
|  | Aq\|pd vs T\|bi | 0.060 | 0.040 | 2.849 | **0.002** |
|  | Aq\|pd vs T\|dd | 0.070 | 0.074 | 1.357 | 0.087 |
|  | Aq\|bi vs SAq\|bi | 0.021 | 0.035 | -0.561 | 0.712 |
|  | Aq\|bi vs T\|bi | 0.027 | 0.046 | -1.245 | 0.884 |
|  | Aq\|bi vs T\|dd | 0.034 | 0.072 | -2.040 | 0.982 |
|  | SAq\|bi vs T\|bi | 0.024 | 0.040 | -0.905 | 0.820 |
|  | SAq\|bi vs T\|dd | 0.029 | 0.070 | -2.305 | 0.990 |
|  | T\|bi vs T\|dd | 0.032 | 0.072 | -2.326 | 0.996 |
| Femur External Shape: Allometry | Aq\|pd vs Aq\|bi | 0.103 | 0.122 | 1.102 | 0.138 |
|  | Aq\|pd vs SAq\|bi | 0.071 | 0.053 | 3.056 | **0.002** |
|  | Aq\|pd vs T\|bi | 0.102 | 0.065 | 3.536 | **0.001** |
|  | Aq\|pd vs T\|dd | 0.063 | 0.055 | 2.250 | **0.008** |
|  | Aq\|bi vs SAq\|bi | 0.079 | 0.122 | -0.034 | 0.501 |
|  | Aq\|bi vs T\|bi | 0.098 | 0.126 | 0.508 | 0.304 |
|  | Aq\|bi vs T\|dd | 0.097 | 0.123 | 0.727 | 0.231 |
|  | SAq\|bi vs T\|bi | 0.072 | 0.074 | 1.488 | 0.077 |
|  | SAq\|bi vs T\|dd | 0.044 | 0.063 | -0.443 | 0.679 |
|  | T\|bi vs T\|dd | 0.075 | 0.073 | 1.766 | **0.041** |
| Humerus Stiffness: Mean  (with sirens) | Aq\|pd vs Aq\|bi | 0.026 | 0.223 | -0.877 | 0.799 |
|  | Aq\|pd vs SAq\|bi | 0.217 | 0.193 | 1.814 | **0.035** |
|  | Aq\|pd vs T\|bi | 0.109 | 0.221 | 0.472 | 0.331 |
|  | Aq\|pd vs T\|dd | 0.307 | 0.188 | 2.647 | **0.002** |
|  | Aq\|bi vs SAq\|bi | 0.190 | 0.176 | 1.701 | **0.031** |
|  | Aq\|bi vs T\|bi | 0.083 | 0.202 | 0.222 | 0.432 |
|  | Aq\|bi vs T\|dd | 0.280 | 0.181 | 2.440 | **0.003** |
|  | SAq\|bi vs T\|bi | 0.108 | 0.167 | 0.888 | 0.205 |
|  | SAq\|bi vs T\|dd | 0.090 | 0.127 | 0.938 | 0.190 |
|  | T\|bi vs T\|dd | 0.198 | 0.166 | 2.011 | **0.021** |
| Humerus Stiffness: Allometry  (with sirens) | Aq\|pd vs Aq\|bi | 0.061 | 0.553 | -0.964 | 0.819 |
|  | Aq\|pd vs SAq\|bi | 0.252 | 0.251 | 1.592 | **0.049** |
|  | Aq\|pd vs T\|bi | 0.100 | 0.348 | -0.153 | 0.572 |
|  | Aq\|pd vs T\|dd | 0.382 | 0.254 | 2.441 | **0.004** |
|  | Aq\|bi vs SAq\|bi | 0.313 | 0.585 | 0.710 | 0.250 |
|  | Aq\|bi vs T\|bi | 0.161 | 0.579 | -0.226 | 0.598 |
|  | Aq\|bi vs T\|dd | 0.443 | 0.536 | 1.182 | 0.120 |
|  | SAq\|bi vs T\|bi | 0.152 | 0.376 | 0.184 | 0.457 |
|  | SAq\|bi vs T\|dd | 0.130 | 0.296 | 0.323 | 0.391 |
|  | T\|bi vs T\|dd | 0.282 | 0.391 | 1.059 | 0.147 |
| Humerus Stiffness: Mean  (no sirens) | Aq\|pd vs Aq\|bi | 0.057 | 0.250 | -0.400 | 0.644 |
|  | Aq\|pd vs SAq\|bi | 0.238 | 0.205 | 1.826 | **0.027** |
|  | Aq\|pd vs T\|bi | 0.135 | 0.227 | 0.734 | 0.260 |
|  | Aq\|pd vs T\|dd | 0.325 | 0.212 | 2.422 | **0.003** |
|  | Aq\|bi vs SAq\|bi | 0.182 | 0.179 | 1.562 | **0.047** |
|  | Aq\|bi vs T\|bi | 0.078 | 0.208 | 0.089 | 0.472 |
|  | Aq\|bi vs T\|dd | 0.268 | 0.183 | 2.301 | **0.004** |
|  | SAq\|bi vs T\|bi | 0.103 | 0.173 | 0.806 | 0.219 |
|  | SAq\|bi vs T\|dd | 0.086 | 0.132 | 0.923 | 0.198 |
|  | T\|bi vs T\|dd | 0.190 | 0.174 | 1.748 | **0.033** |
| Femur Stiffness: Mean | Aq\|pd vs Aq\|bi | 0.256 | 0.275 | 1.425 | 0.071 |
|  | Aq\|pd vs SAq\|bi | 0.300 | 0.231 | 2.099 | **0.015** |
|  | Aq\|pd vs T\|bi | 0.256 | 0.260 | 1.562 | 0.058 |
|  | Aq\|pd vs T\|dd | 0.239 | 0.237 | 1.604 | **0.048** |
|  | Aq\|bi vs SAq\|bi | 0.044 | 0.222 | -0.472 | 0.686 |
|  | Aq\|bi vs T\|bi | 0.000 | 0.230 | -2.440 | 0.999 |
|  | Aq\|bi vs T\|dd | 0.017 | 0.209 | -1.300 | 0.890 |
|  | SAq\|bi vs T\|bi | 0.045 | 0.185 | -0.321 | 0.625 |
|  | SAq\|bi vs T\|dd | 0.062 | 0.141 | 0.267 | 0.405 |
|  | T\|bi vs T\|dd | 0.017 | 0.184 | -1.072 | 0.846 |
| Humerus Density: Mean  (with sirens) | Aq\|pd vs Aq\|bi | 0.014 | 0.066 | -0.411 | 0.641 |
|  | Aq\|pd vs SAq\|bi | 0.020 | 0.058 | 0.014 | 0.505 |
|  | Aq\|pd vs T\|bi | 0.029 | 0.059 | 0.379 | 0.367 |
|  | Aq\|pd vs T\|dd | 0.088 | 0.056 | 2.522 | **0.002** |
|  | Aq\|bi vs SAq\|bi | 0.034 | 0.052 | 0.851 | 0.211 |
|  | Aq\|bi vs T\|bi | 0.043 | 0.059 | 1.092 | 0.138 |
|  | Aq\|bi vs T\|dd | 0.102 | 0.050 | 3.040 | **0.001** |
|  | SAq\|bi vs T\|bi | 0.009 | 0.047 | -0.634 | 0.726 |
|  | SAq\|bi vs T\|dd | 0.068 | 0.039 | 2.726 | **0.001** |
|  | T\|bi vs T\|dd | 0.059 | 0.047 | 2.111 | **0.011** |
| Humerus Density: Allometry  (with sirens) | Aq\|pd vs Aq\|bi | 0.009 | 0.151 | -1.429 | 0.919 |
|  | Aq\|pd vs SAq\|bi | 0.039 | 0.072 | 0.637 | 0.281 |
|  | Aq\|pd vs T\|bi | 0.021 | 0.105 | -0.529 | 0.700 |
|  | Aq\|pd vs T\|dd | 0.134 | 0.072 | 2.922 | **0.002** |
|  | Aq\|bi vs SAq\|bi | 0.030 | 0.164 | -0.525 | 0.695 |
|  | Aq\|bi vs T\|bi | 0.012 | 0.173 | -1.302 | 0.884 |
|  | Aq\|bi vs T\|dd | 0.125 | 0.159 | 1.188 | 0.126 |
|  | SAq\|bi vs T\|bi | 0.018 | 0.110 | -0.760 | 0.766 |
|  | SAq\|bi vs T\|dd | 0.095 | 0.086 | 1.737 | **0.031** |
|  | T\|bi vs T\|dd | 0.113 | 0.110 | 1.616 | **0.045** |
| Humerus Density: Mean  (no sirens) | Aq\|pd vs Aq\|bi | 0.000 | 0.073 | -2.236 | 0.989 |
|  | Aq\|pd vs SAq\|bi | 0.034 | 0.061 | 0.608 | 0.301 |
|  | Aq\|pd vs T\|bi | 0.043 | 0.069 | 0.854 | 0.207 |
|  | Aq\|pd vs T\|dd | 0.099 | 0.062 | 2.450 | **0.002** |
|  | Aq\|bi vs SAq\|bi | 0.033 | 0.053 | 0.805 | 0.213 |
|  | Aq\|bi vs T\|bi | 0.043 | 0.060 | 1.033 | 0.156 |
|  | Aq\|bi vs T\|dd | 0.099 | 0.054 | 2.895 | **0.001** |
|  | SAq\|bi vs T\|bi | 0.009 | 0.049 | -0.558 | 0.689 |
|  | SAq\|bi vs T\|dd | 0.065 | 0.037 | 2.666 | **0.002** |
|  | T\|bi vs T\|dd | 0.056 | 0.050 | 1.797 | **0.029** |
| Humerus Density: Allometry  (no sirens) | Aq\|pd vs Aq\|bi | 0.002 | 0.157 | -2.071 | 0.980 |
|  | Aq\|pd vs SAq\|bi | 0.028 | 0.080 | 0.042 | 0.510 |
|  | Aq\|pd vs T\|bi | 0.010 | 0.103 | -0.979 | 0.817 |
|  | Aq\|pd vs T\|dd | 0.123 | 0.075 | 2.524 | **0.002** |
|  | Aq\|bi vs SAq\|bi | 0.030 | 0.166 | -0.555 | 0.708 |
|  | Aq\|bi vs T\|bi | 0.012 | 0.174 | -1.272 | 0.884 |
|  | Aq\|bi vs T\|dd | 0.125 | 0.157 | 1.181 | 0.127 |
|  | SAq\|bi vs T\|bi | 0.018 | 0.114 | -0.683 | 0.744 |
|  | SAq\|bi vs T\|dd | 0.095 | 0.089 | 1.693 | **0.031** |
|  | T\|bi vs T\|dd | 0.113 | 0.110 | 1.608 | **0.039** |
| Femur Density: Mean | Aq\|pd vs Aq\|bi | 0.038 | 0.072 | 0.506 | 0.326 |
|  | Aq\|pd vs SAq\|bi | 0.041 | 0.065 | 0.845 | 0.223 |
|  | Aq\|pd vs T\|bi | 0.062 | 0.069 | 1.420 | 0.076 |
|  | Aq\|pd vs T\|dd | 0.089 | 0.065 | 2.246 | **0.009** |
|  | Aq\|bi vs SAq\|bi | 0.004 | 0.054 | -1.419 | 0.903 |
|  | Aq\|bi vs T\|bi | 0.024 | 0.062 | 0.228 | 0.425 |
|  | Aq\|bi vs T\|dd | 0.052 | 0.053 | 1.511 | 0.057 |
|  | SAq\|bi vs T\|bi | 0.021 | 0.049 | 0.326 | 0.393 |
|  | SAq\|bi vs T\|dd | 0.048 | 0.038 | 2.041 | **0.010** |
|  | T\|bi vs T\|dd | 0.027 | 0.051 | 0.658 | 0.271 |

**Table S5.** Results of the phylogenetic paired t-tests comparing the stiffness and density of the forelimbs and hindlimbs. Bold values indicate statistical significance (p < 0.05). A = Aquatic, SAq = Semi-aquatic, and T = Terrestrial; pd = Paedomorphic, bi = Multiphasic, and dd = Direct developer.

|  | **Stiffness** | | | **Density** | | |
| --- | --- | --- | --- | --- | --- | --- |
|  | **Df** | **t** | **p-value** | **Df** | **t** | **p-value** |
| Aq\|pd | 10 | -0.869 | 0.405 | 10 | 0.196 | 0.848 |
| Aq\|bi | 9 | -0.787 | 0.451 | 9 | 3.287 | **0.009** |
| SAq\|bi | 32 | 0.608 | 0.547 | 32 | -0.677 | 0.503 |
| T\|bi | 16 | -1.247 | 0.230 | 16 | 1.192 | 0.251 |
| T\|dd | 47 | 0.873 | 0.387 | 47 | -0.330 | 0.743 |

**Table S6.** Evolutionary rates of external limb shape by ecotype. Humeral analyses were performed with and without the sirens included in the dataset. A = Aquatic, SAq = Semi-aquatic, and T = Terrestrial; pd = Paedomorphic, bi = Multiphasic, and dd = Direct developer.

|  | **Aq\|pd** | **Aq\|bi** | **Saq\|bi** | **T\|bi** | **T\|dd** |
| --- | --- | --- | --- | --- | --- |
| Humerus Shape (with sirens) | 6.48E-07 | 1.91E-07 | 2.60E-07 | 3.94E-07 | 1.16E-07 |
| Humerus Shape (no sirens) | 7.29E-07 | 1.92E-07 | 2.59E-07 | 3.92E-07 | 1.16E-07 |
| Femur Shape | 7.04E-07 | 1.65E-07 | 2.35E-07 | 4.00E-07 | 1.33E-07 |

**Table S7.** Fit of the 26 *hOUwie* models used to model the evolution of humeral stiffness with sirens. Model names reflect the different parameters being varied. Models were fit with either a Brownian Motion (“bm”) or an Ornstein-Uhlenbeck (“ou”) model of evolution with rate (σ^2^, “v”) and optima (θ, “m”) parameters that vary with different discrete character states. We used three different regime classifications (“ecotype”, “habitat”, lifecycle”). Character dependent models (“cd”) indicate that variation in parameter values is attributable to the character state themselves, whereas character independent models (“cid”) indicate that other factors explain the variation in parameters, as represented by hidden states in the model. For instance, the best fitting model “cd_oumv_habitat” indicates that humeral stiffness is described by an Ornstein-Uhlenbeck model of evolution and that aquatic, semi-aquatic, and terrestrial species are evolving at different rates towards different optima. That contrasts “cd_oumv_lifecycle”, where the rates and optima differ between life cycle strategies (i.e., paedomorphic, biphasic, and direct developer). We used all models and their weighted AICc scores to calculate the model averaged rate and optima parameters for each ecotype. The best fitting models, as determined by a Δ2.0 AICc cutoff, are bolded.

| **Model** | **np** | **lnLik** | **DiscLik** | **ContLik** | **AICc** | **dAICc** | **AICcwt** |
| --- | --- | --- | --- | --- | --- | --- | --- |
| cid_bm1 | 14 | -140.29 | -136.66 | -1.38 | 312.13 | 37.93 | 3.23E-09 |
| cid_ou1 | 15 | -131.66 | -136.16 | 5.81 | 297.42 | 23.23 | 5.05E-06 |
| cd_bmv_ecotype | 11 | -135.86 | -139.44 | 5.63 | 295.91 | 21.71 | 1.08E-05 |
| **cd_ouv_ecotype** | **12** | **-124.04** | **-139.75** | **13.14** | **274.69** | **0.49** | **4.36E-01** |
| cd_oum_ecotype | 12 | -132.21 | -138.45 | 4.76 | 291.02 | 16.82 | 1.24E-04 |
| cd_oumv_ecotype | 16 | -124.27 | -141.54 | 15.66 | 285.23 | 11.03 | 2.24E-03 |
| cid_bmv_ecotype | 15 | -140.40 | -134.94 | -1.34 | 314.91 | 40.71 | 8.06E-10 |
| cid_ouv_ecotype | 16 | -131.88 | -134.20 | 5.23 | 300.45 | 26.25 | 1.11E-06 |
| cid_oum_ecotype | 16 | -136.77 | -135.04 | 1.38 | 310.22 | 36.03 | 8.39E-09 |
| cid_oumv_ecotype | 17 | -138.07 | -134.86 | 1.86 | 315.45 | 41.26 | 6.14E-10 |
| cd_bmv_habitat | 9 | -136.39 | -140.82 | 4.58 | 292.24 | 18.05 | 6.73E-05 |
| cd_ouv_habitat | 11 | -133.69 | -139.61 | 6.04 | 291.56 | 17.36 | 9.47E-05 |
| cd_oum_habitat | 10 | -133.13 | -141.57 | 7.70 | 288.06 | 13.86 | 5.45E-04 |
| **cd_oumv_habitat** | **12** | **-123.80** | **-139.65** | **14.71** | **274.20** | **0.00** | **5.58E-01** |
| cid_bmv_habitat | 19 | -130.73 | -134.65 | 9.33 | 306.20 | 32.00 | 6.29E-08 |
| cid_ouv_habitat | 24 | -121.06 | -136.71 | 19.81 | 301.24 | 27.04 | 7.49E-07 |
| cid_oum_habitat | 20 | -131.34 | -138.02 | 8.55 | 310.18 | 35.99 | 8.56E-09 |
| cid_oumv_habitat | 25 | -120.46 | -136.42 | 19.13 | 303.07 | 28.87 | 3.00E-07 |
| cd_bmv_lifecycle | 9 | -135.46 | -141.02 | 3.12 | 290.38 | 16.18 | 1.71E-04 |
| cd_ouv_lifecycle | 11 | -138.60 | -141.86 | 1.62 | 301.39 | 27.19 | 6.95E-07 |
| cd_oum_lifecycle | 10 | -134.84 | -141.93 | 6.42 | 291.49 | 17.29 | 9.83E-05 |
| cd_oumv_lifecycle | 12 | -129.33 | -140.53 | 11.22 | 285.26 | 11.06 | 2.21E-03 |
| cid_bmv_lifecycle | 19 | -135.15 | -136.08 | 4.15 | 315.02 | 40.82 | 7.63E-10 |
| cid_ouv_lifecycle | 24 | -125.27 | -135.52 | 13.89 | 309.64 | 35.44 | 1.12E-08 |
| cid_oum_lifecycle | 20 | -139.09 | -134.39 | -1.17 | 325.68 | 51.49 | 3.69E-12 |
| cid_oumv_lifecycle | 25 | -122.31 | -131.42 | 11.06 | 306.78 | 32.58 | 4.70E-08 |

**Table S8.** Fit of the 26 *hOUwie* models used to model the evolution of humeral stiffness without sirens. Interpretation of the name and structures are the same as Table S7. The best fitting models, as determined by a Δ2.0 AICc cutoff, are bolded.

| **Model** | **np** | **lnLik** | **DiscLik** | **ContLik** | **AICc** | **dAICc** | **AICcwt** |
| --- | --- | --- | --- | --- | --- | --- | --- |
| cid_bm1 | 14 | -176.70 | -176.84 | -6.82 | 385.09 | 124.94 | 7.38E-28 |
| cid_ou1 | 15 | -129.20 | -128.24 | 3.15 | 292.64 | 32.49 | 8.78E-08 |
| cd_bmv_ecotype | 11 | -134.41 | -136.08 | 3.11 | 293.08 | 32.93 | 7.06E-08 |
| cd_ouv_ecotype | 12 | -131.96 | -140.55 | 7.78 | 290.60 | 30.45 | 2.44E-07 |
| cd_oum_ecotype | 12 | -138.04 | -138.87 | 1.02 | 302.76 | 42.61 | 5.57E-10 |
| cd_oumv_ecotype | 16 | -123.89 | -137.97 | 12.98 | 284.63 | 24.48 | 4.83E-06 |
| cid_bmv_ecotype | 15 | -135.18 | -129.29 | -2.27 | 304.61 | 44.46 | 2.22E-10 |
| cid_ouv_ecotype | 16 | -137.15 | -135.24 | 0.04 | 311.16 | 51.02 | 8.34E-12 |
| cid_oum_ecotype | 16 | -136.95 | -133.38 | -0.44 | 310.76 | 50.61 | 1.02E-11 |
| cid_oumv_ecotype | 17 | -137.57 | -132.73 | -1.30 | 314.66 | 54.51 | 1.45E-12 |
| cd_bmv_habitat | 9 | -132.29 | -138.16 | 6.59 | 284.10 | 23.95 | 6.28E-06 |
| cd_ouv_habitat | 11 | -128.73 | -129.75 | 3.93 | 281.72 | 21.57 | 2.07E-05 |
| cd_oum_habitat | 10 | -136.38 | -139.00 | -0.09 | 294.63 | 34.49 | 3.24E-08 |
| **cd_oumv_habitat** | **12** | **-116.73** | **-131.34** | **16.47** | **260.15** | **0.00** | **9.98E-01** |
| cid_bmv_habitat | 19 | -128.61 | -132.61 | 9.77 | 302.20 | 42.05 | 7.39E-10 |
| cid_ouv_habitat | 24 | -133.32 | -139.92 | 8.48 | 326.18 | 66.03 | 4.57E-15 |
| cid_oum_habitat | 20 | -132.69 | -140.31 | 8.56 | 313.16 | 53.01 | 3.08E-12 |
| cid_oumv_habitat | 25 | -115.48 | -132.45 | 21.18 | 293.57 | 33.43 | 5.51E-08 |
| cd_bmv_lifecycle | 9 | -132.77 | -129.75 | 0.98 | 285.05 | 24.90 | 3.91E-06 |
| cd_ouv_lifecycle | 11 | -135.39 | -135.99 | 1.18 | 295.03 | 34.88 | 2.66E-08 |
| cd_oum_lifecycle | 10 | -125.65 | -128.42 | 6.08 | 273.17 | 13.03 | 1.48E-03 |
| cd_oumv_lifecycle | 12 | -129.14 | -134.86 | 5.32 | 284.97 | 24.82 | 4.08E-06 |
| cid_bmv_lifecycle | 19 | -129.90 | -133.05 | 6.40 | 304.77 | 44.62 | 2.04E-10 |
| cid_ouv_lifecycle | 24 | -122.78 | -128.91 | 9.82 | 305.10 | 44.96 | 1.73E-10 |
| cid_oum_lifecycle | 20 | -134.41 | -130.80 | -0.77 | 316.59 | 56.44 | 5.54E-13 |
| cid_oumv_lifecycle | 25 | -120.43 | -129.00 | 12.52 | 303.48 | 43.33 | 3.89E-10 |

**Table S9.** Fit of the 26 *hOUwie* models used to model the evolution of femoral stiffness. Interpretation of the name and structures are the same as Table S7. The best fitting models, as determined by a Δ2.0 AICc cutoff, are bolded.

| **Model** | **np** | **lnLik** | **DiscLik** | **ContLik** | **AICc** | **dAICc** | **AICcwt** |
| --- | --- | --- | --- | --- | --- | --- | --- |
| cid_bm1 | 14 | -164.26 | -129.70 | -32.10 | 360.21 | 36.26 | 7.57E-09 |
| cid_ou1 | 15 | -158.27 | -137.51 | -18.60 | 350.78 | 26.83 | 8.45E-07 |
| cd_bmv_ecotype | 11 | -173.55 | -139.01 | -35.51 | 371.36 | 47.40 | 2.87E-11 |
| cd_ouv_ecotype | 12 | -149.99 | -129.50 | -17.54 | 326.68 | 2.72 | 1.45E-01 |
| cd_oum_ecotype | 12 | -156.74 | -132.08 | -25.80 | 340.16 | 16.21 | 1.70E-04 |
| cd_oumv_ecotype | 16 | -154.87 | -128.79 | -22.23 | 346.60 | 22.65 | 6.82E-06 |
| cid_bmv_ecotype | 15 | -168.17 | -136.62 | -31.75 | 370.59 | 46.64 | 4.22E-11 |
| cid_ouv_ecotype | 16 | -151.65 | -129.97 | -19.75 | 340.16 | 16.20 | 1.71E-04 |
| cid_oum_ecotype | 16 | -151.55 | -130.51 | -18.28 | 339.95 | 16.00 | 1.90E-04 |
| cid_oumv_ecotype | 17 | -154.30 | -133.27 | -17.99 | 348.12 | 24.17 | 3.20E-06 |
| cd_bmv_habitat | 9 | -166.71 | -133.11 | -32.83 | 352.92 | 28.97 | 2.89E-07 |
| cd_ouv_habitat | 11 | -168.85 | -141.84 | -29.05 | 361.96 | 38.01 | 3.15E-09 |
| cd_oum_habitat | 10 | -152.58 | -133.78 | -16.39 | 327.03 | 3.08 | 1.21E-01 |
| **cd_oumv_habitat** | **12** | **-148.63** | **-130.28** | **-16.66** | **323.95** | **0.00** | **5.65E-01** |
| cid_bmv_habitat | 19 | -165.66 | -130.92 | -31.75 | 376.30 | 52.35 | 2.42E-12 |
| cid_ouv_habitat | 24 | -167.37 | -135.67 | -28.84 | 394.28 | 70.33 | 3.03E-16 |
| cid_oum_habitat | 20 | -159.90 | -127.85 | -26.10 | 367.57 | 43.62 | 1.91E-10 |
| cid_oumv_habitat | 25 | -153.79 | -134.64 | -17.61 | 370.20 | 46.25 | 5.11E-11 |
| cd_bmv_lifecycle | 9 | -166.21 | -133.32 | -31.61 | 351.93 | 27.98 | 4.76E-07 |
| cd_ouv_lifecycle | 11 | -167.38 | -134.37 | -30.78 | 359.02 | 35.07 | 1.37E-08 |
| cd_oum_lifecycle | 10 | -152.26 | -127.93 | -20.87 | 326.37 | 2.42 | 1.68E-01 |
| cd_oumv_lifecycle | 12 | -160.79 | -139.31 | -21.82 | 348.26 | 24.31 | 2.97E-06 |
| cid_bmv_lifecycle | 19 | -178.87 | -133.66 | -43.65 | 402.72 | 78.77 | 4.45E-18 |
| cid_ouv_lifecycle | 24 | -147.86 | -130.48 | -15.44 | 355.26 | 31.31 | 8.98E-08 |
| cid_oum_lifecycle | 20 | -156.63 | -133.97 | -19.23 | 361.04 | 37.09 | 4.99E-09 |
| cid_oumv_lifecycle | 25 | -148.78 | -128.59 | -16.54 | 360.18 | 36.23 | 7.67E-09 |

**Table S10.** Fit of the 26 *hOUwie* models used to model the evolution of humeral density with sirens. Interpretation of the name and structures are the same as Table S7. The best fitting models, as determined by a Δ2.0 AICc cutoff, are bolded.

| **Model** | **np** | **lnLik** | **DiscLik** | **ContLik** | **AICc** | **dAICc** | **AICcwt** |
| --- | --- | --- | --- | --- | --- | --- | --- |
| cid_bm1 | 14 | 31.62 | -136.63 | 170.43 | -31.68 | 21.21 | 2.03E-05 |
| cid_ou1 | 15 | 40.38 | -133.77 | 177.01 | -46.66 | 6.23 | 3.64E-02 |
| cd_bmv_ecotype | 11 | 32.19 | -144.02 | 166.62 | -40.21 | 12.68 | 1.44E-03 |
| cd_ouv_ecotype | 12 | 19.07 | -137.63 | 158.23 | -11.54 | 41.35 | 8.59E-10 |
| cd_oum_ecotype | 12 | 25.61 | -141.13 | 164.42 | -24.61 | 28.28 | 5.93E-07 |
| cd_oumv_ecotype | 16 | 42.28 | -139.15 | 182.12 | -47.88 | 5.02 | 6.68E-02 |
| cid_bmv_ecotype | 15 | 33.27 | -134.49 | 170.71 | -32.43 | 20.46 | 2.96E-05 |
| cid_ouv_ecotype | 16 | 39.32 | -133.55 | 175.51 | -41.95 | 10.94 | 3.46E-03 |
| cid_oum_ecotype | 16 | 36.07 | -134.50 | 174.81 | -35.45 | 17.45 | 1.34E-04 |
| cid_oumv_ecotype | 17 | 36.78 | -135.79 | 174.67 | -34.23 | 18.66 | 7.28E-05 |
| cd_bmv_habitat | 9 | 32.88 | -138.50 | 174.33 | -46.30 | 6.60 | 3.03E-02 |
| cd_ouv_habitat | 11 | -71.84 | -142.37 | 68.89 | 167.87 | 220.76 | 9.47E-49 |
| cd_oum_habitat | 10 | 26.42 | -142.12 | 168.95 | -31.03 | 21.86 | 1.47E-05 |
| **cd_oumv_habitat** | **12** | **39.75** | **-138.23** | **180.46** | **-52.89** | **0.00** | **8.20E-01** |
| cid_bmv_habitat | 19 | 12.66 | -157.77 | 167.96 | 19.41 | 72.30 | 1.63E-16 |
| cid_ouv_habitat | 24 | 32.25 | -133.90 | 170.05 | -5.39 | 47.51 | 3.96E-11 |
| cid_oum_habitat | 20 | 35.53 | -137.39 | 175.11 | -23.55 | 29.34 | 3.49E-07 |
| cid_oumv_habitat | 25 | 36.85 | -135.51 | 175.19 | -11.54 | 41.35 | 8.62E-10 |
| cd_bmv_lifecycle | 9 | 33.15 | -139.86 | 172.58 | -46.83 | 6.06 | 3.97E-02 |
| cd_ouv_lifecycle | 11 | 31.64 | -140.72 | 172.04 | -39.10 | 13.79 | 8.30E-04 |
| cd_oum_lifecycle | 10 | 30.02 | -139.56 | 170.84 | -38.24 | 14.65 | 5.39E-04 |
| cd_oumv_lifecycle | 12 | 29.07 | -140.25 | 168.73 | -31.55 | 21.34 | 1.90E-05 |
| cid_bmv_lifecycle | 19 | 31.86 | -140.96 | 175.27 | -18.99 | 33.90 | 3.57E-08 |
| cid_ouv_lifecycle | 24 | 31.09 | -134.54 | 168.89 | -3.06 | 49.83 | 1.24E-11 |
| cid_oum_lifecycle | 20 | 36.59 | -135.84 | 174.72 | -25.68 | 27.21 | 1.01E-06 |
| cid_oumv_lifecycle | 25 | 29.59 | -135.99 | 167.11 | 2.97 | 55.86 | 6.09E-13 |

**Table S11.** Fit of the 26 *hOUwie* models used to model the evolution of humeral density without sirens. Interpretation of the name and structures are the same as Table S7. The best fitting models, as determined by a Δ2.0 AICc cutoff, are bolded.

| **Model** | **np** | **lnLik** | **DiscLik** | **ContLik** | **AICc** | **dAICc** | **AICcwt** |
| --- | --- | --- | --- | --- | --- | --- | --- |
| cid_bm1 | 14 | 31.40 | -134.02 | 164.61 | -31.12 | 27.96 | 4.43E-07 |
| cid_ou1 | 15 | 39.62 | -129.34 | 171.35 | -44.98 | 14.09 | 4.53E-04 |
| cd_bmv_ecotype | 11 | 16.63 | -139.99 | 157.97 | -8.99 | 50.08 | 6.94E-12 |
| cd_ouv_ecotype | 12 | -3.33 | -139.01 | 133.76 | 33.34 | 92.42 | 4.44E-21 |
| cd_oum_ecotype | 12 | 17.83 | -138.55 | 157.47 | -8.98 | 50.10 | 6.88E-12 |
| cd_oumv_ecotype | 16 | 41.17 | -133.57 | 177.83 | -45.48 | 13.59 | 5.82E-04 |
| cid_bmv_ecotype | 15 | 31.91 | -131.73 | 165.74 | -29.56 | 29.51 | 2.03E-07 |
| cid_ouv_ecotype | 16 | 15.77 | -128.89 | 148.13 | 5.31 | 64.39 | 5.44E-15 |
| cid_oum_ecotype | 16 | 29.02 | -134.56 | 166.01 | -21.19 | 37.89 | 3.08E-09 |
| cid_oumv_ecotype | 17 | 32.71 | -133.18 | 169.39 | -25.90 | 33.18 | 3.25E-08 |
| cd_bmv_habitat | 9 | 35.34 | -132.20 | 171.07 | -51.17 | 7.91 | 9.98E-03 |
| cd_ouv_habitat | 11 | -47.75 | -139.35 | 89.96 | 119.77 | 178.84 | 7.61E-40 |
| cd_oum_habitat | 10 | 15.08 | -138.84 | 155.38 | -8.29 | 50.79 | 4.88E-12 |
| cd_oumv_habitat | 12 | 18.37 | -138.94 | 154.94 | -10.05 | 49.03 | 1.18E-11 |
| cid_bmv_habitat | 19 | 32.96 | -132.73 | 170.17 | -20.95 | 38.13 | 2.73E-09 |
| cid_ouv_habitat | 24 | -38.43 | -139.10 | 104.49 | 136.40 | 195.48 | 1.86E-43 |
| cid_oum_habitat | 20 | 35.92 | -133.71 | 172.19 | -24.07 | 35.01 | 1.30E-08 |
| cid_oumv_habitat | 25 | 48.74 | -144.45 | 200.33 | -34.86 | 24.22 | 2.87E-06 |
| cd_bmv_lifecycle | 9 | 28.12 | -134.48 | 165.99 | -36.72 | 22.36 | 7.28E-06 |
| cd_ouv_lifecycle | 11 | 31.76 | -136.42 | 166.69 | -39.26 | 19.82 | 2.59E-05 |
| **cd_oum_lifecycle** | **10** | **40.36** | **-129.10** | **172.31** | **-58.86** | **0.22** | **4.67E-01** |
| **cd_oumv_lifecycle** | **12** | **42.88** | **-128.39** | **174.50** | **-59.08** | **0.00** | **5.21E-01** |
| cid_bmv_lifecycle | 19 | 46.27 | -131.78 | 179.74 | -47.56 | 11.52 | 1.64E-03 |
| cid_ouv_lifecycle | 24 | 34.69 | -127.61 | 166.56 | -9.85 | 49.23 | 1.06E-11 |
| cid_oum_lifecycle | 20 | 36.64 | -130.65 | 170.39 | -25.51 | 33.57 | 2.67E-08 |
| cid_oumv_lifecycle | 25 | 32.33 | -127.85 | 163.74 | -2.03 | 57.05 | 2.13E-13 |

**Table S12.** Fit of the 26 *hOUwie* models used to model the evolution of femoral density. Interpretation of the name and structures are the same as Table S7. The best fitting models, as determined by a Δ2.0 AICc cutoff, are bolded.

| **Model** | **np** | **lnLik** | **DiscLik** | **ContLik** | **AICc** | **dAICc** | **AICcwt** |
| --- | --- | --- | --- | --- | --- | --- | --- |
| cid_bm1 | 14 | 3.24 | -128.88 | 135.55 | 25.20 | 33.12 | 3.53E-08 |
| cid_ou1 | 15 | 18.09 | -128.68 | 150.00 | -1.93 | 5.99 | 2.75E-02 |
| cd_bmv_ecotype | 11 | 12.07 | -129.63 | 143.10 | 0.12 | 8.05 | 9.83E-03 |
| **cd_ouv_ecotype** | **12** | **16.88** | **-129.99** | **149.07** | **-7.07** | **0.85** | **3.58E-01** |
| cd_oum_ecotype | 12 | 10.19 | -140.66 | 149.16 | 6.31 | 14.23 | 4.47E-04 |
| cd_oumv_ecotype | 16 | 13.55 | -138.23 | 153.31 | 9.75 | 17.67 | 7.98E-05 |
| cid_bmv_ecotype | 15 | 3.78 | -128.86 | 136.01 | 26.68 | 34.60 | 1.68E-08 |
| cid_ouv_ecotype | 16 | 14.35 | -128.29 | 146.66 | 8.15 | 16.07 | 1.78E-04 |
| cid_oum_ecotype | 16 | 19.06 | -129.97 | 151.18 | -1.27 | 6.66 | 1.97E-02 |
| cid_oumv_ecotype | 17 | 9.67 | -135.46 | 146.52 | 20.17 | 28.10 | 4.35E-07 |
| cd_bmv_habitat | 9 | 2.60 | -131.08 | 136.55 | 14.31 | 22.23 | 8.17E-06 |
| cd_ouv_habitat | 11 | -37.26 | -132.88 | 95.23 | 98.78 | 106.71 | 3.70E-24 |
| **cd_oum_habitat** | **10** | **14.89** | **-134.42** | **151.09** | **-7.92** | **0.00** | **5.49E-01** |
| cd_oumv_habitat | 12 | -2.59 | -132.93 | 133.30 | 31.87 | 39.79 | 1.26E-09 |
| cid_bmv_habitat | 19 | 3.75 | -132.86 | 139.11 | 37.48 | 45.40 | 7.62E-11 |
| cid_ouv_habitat | 24 | 21.05 | -129.83 | 153.34 | 17.44 | 25.37 | 1.70E-06 |
| cid_oum_habitat | 20 | 22.07 | -127.30 | 152.98 | 3.63 | 11.55 | 1.70E-03 |
| cid_oumv_habitat | 25 | 17.90 | -132.07 | 154.25 | 26.83 | 34.75 | 1.56E-08 |
| cd_bmv_lifecycle | 9 | -1.30 | -136.69 | 137.01 | 22.12 | 30.04 | 1.65E-07 |
| cd_ouv_lifecycle | 11 | -1.13 | -135.13 | 135.02 | 26.52 | 34.44 | 1.83E-08 |
| cd_oum_lifecycle | 10 | 12.08 | -139.05 | 152.20 | -2.29 | 5.63 | 3.29E-02 |
| cd_oumv_lifecycle | 12 | -1.28 | -138.79 | 138.03 | 29.24 | 37.16 | 4.67E-09 |
| cid_bmv_lifecycle | 19 | 2.63 | -132.64 | 137.84 | 39.71 | 47.63 | 2.50E-11 |
| cid_ouv_lifecycle | 24 | 21.97 | -137.00 | 163.25 | 15.60 | 23.52 | 4.28E-06 |
| cid_oum_lifecycle | 20 | 11.78 | -131.30 | 145.53 | 24.22 | 32.14 | 5.77E-08 |
| cid_oumv_lifecycle | 25 | 5.11 | -143.23 | 155.03 | 52.40 | 60.33 | 4.37E-14 |

**Table S13.** Model averaged evolutionary rates and optima of each cross-sectional trait across ecotypes. Parameters for each trait were estimated with 26 *hOUwie* and averaged with the weighted AICc scores (Tables S7-12). Aq = Aquatic, SAq = Semi-Aquatic, T = Terrestrial; pd = Paedomorphic, bi = multiphasic, and dd = Direct developer.

|  | **Trait** | **Aq\|pd** | **Aq\|bi** | **SAq\|bi** | **T\|bi** | **T\|dd** |
| --- | --- | --- | --- | --- | --- | --- |
| Evolutionary Rates | Humeral Stiffness (with sirens) | 4.56E-04 | 2.81E-04 | 4.49E-03 | 2.75E-03 | 3.69E-03 |
|  | Humeral Stiffness (no sirens) | 3.64E-04 | 3.64E-04 | 3.02E-03 | 2.89E-03 | 2.89E-03 |
|  | Femoral Stiffness | 2.86E-03 | 2.51E-03 | 4.23E-03 | 3.51E-03 | 3.34E-03 |
|  | Humeral Density (with sirens) | 8.81E-05 | 8.61E-05 | 2.05E-04 | 1.45E-04 | 1.53E-04 |
|  | Humeral Density (no sirens) | 1.11E-04 | 2.03E-04 | 2.04E-04 | 2.03E-04 | 2.26E-04 |
|  | Femoral Density | 4.80E-04 | 3.77E-04 | 3.90E-04 | 5.75E-04 | 7.58E-04 |
| Trait Optima | Humeral Stiffness (with sirens) | 1.265 | 1.265 | 1.444 | 1.380 | 1.380 |
|  | Humeral Stiffness (no sirens) | 1.136 | 1.136 | 1.606 | 1.427 | 1.427 |
|  | Femoral Stiffness | 1.150 | 1.313 | 1.398 | 1.434 | 1.423 |
|  | Humeral Density (with sirens) | 0.955 | 0.955 | 0.941 | 0.921 | 0.926 |
|  | Humeral Density (no sirens) | 0.980 | 0.935 | 0.935 | 0.935 | 0.909 |
|  | Femoral Density | 0.936 | 0.935 | 0.940 | 0.921 | 0.920 |
